# An animal component free bioprocess to synthesize 3D human matrix scaffolds using mesenchymal stromal cells

**DOI:** 10.1101/2025.04.17.649452

**Authors:** Shaianne N. Stein, Lorena R. Braid

## Abstract

Basement membrane is a specialized extracellular matrix (ECM) that compartmentalizes epithelial and endothelial tissues and provides essential structural and signaling cues for tissue organization. While fibrillar collagens (Col) of the interstitial matrix (e.g., Col I and Col III) are widely used in tissue modeling, the networking collagens that scaffold basement membrane, including human Col IV and Col VI, remain difficult to access. Commercial basement membrane surrogates such as Matrigel® are derived from murine tumors and are ill-defined, dilute, variable, and incompatible with animal-free biomanufacturing. Thus, there is a critical need for human-derived basement membrane matrices that are free of xenogenic contaminants and do not rely on animal agriculture.

Here, we investigated whether human mesenchymal stromal cells (MSCs) could serve as a platform to produce self-assembling basement membrane components under chemically defined, xeno-free conditions. MSCs from placental, umbilical cord, bone marrow, and adipose tissues were cultured as three-dimensional spheroids and adherent multilayered sheets. Confocal imaging of whole-mount, decellularized spheroid matrices revealed complex networks of fibronectin and Col IV with topological and organizational features characteristic of native basement membrane. Perinatal MSCs produced distinct matrix architectures consisting of apical fibronectin sheets underlaid by continuous Col IV networks. Time-resolved imaging of umbilical cord MSC–derived matrix sheets demonstrated a reproducible sequence of basement membrane assembly that parallels developmental tissue organization.

Together, these findings demonstrate that human MSCs, cultured entirely without animal-derived components, can synthesize functional basement membrane proteins that self-assemble into ordered, tissue-like scaffolds. This work establishes MSCs as a scalable, sustainable, and cruelty-free platform for the manufacturing of human basement membrane matrices for bioengineering and regenerative medicine applications.

## 1 Introduction

The extracellular matrix (ECM) plays a central role in tissue organization, morphogenesis, and cell fate determination, making it a critical component of bioengineered human tissues and organoid systems (Jain et al., 2022; Kozlowski et al., 2021; Motta et al., 2023; Olesen et al., 2021). While interstitial matrix provides essential structural support, tensile strength, and scaffolding for cells within connective tissues, basement membrane–type matrices provide essential biochemical and structural cues that regulate epithelial and endothelial polarity, vascularization, and growth factor signaling (Jain et al., 2022; Jayadev & Sherwood, 2017; Khalilgharibi & Mao, 2021; Pompili et al., 2021; Sekiguchi & Yamada, 2018; Yurchenco, 2011). Despite their importance, the development of human-relevant basement membrane scaffolds remains limited by the availability of appropriate matrix materials (Aisenbrey & Murphy, 2020; Perugini & Santin, 2021).

Currently, basement membrane surrogates such as Matrigel® and Cultrex™ are widely used in three-dimensional (3D) culture systems (Aisenbrey & Murphy, 2020; Kane et al., 2018). However, these materials are derived from murine Engelbreth–Holm–Swarm sarcoma and only distantly resemble native human basement membrane in composition and organization (Huang et al., 2021; Hughes et al., 2010; Kleinman & Martin, 2005; Kozlowski et al., 2021). Their reliance on animal agriculture raises ethical and sustainability concerns, while xenogeneic contaminants, batch-to-batch variability, and undefined bioactive components limit experimental reproducibility and translational applicability (Aisenbrey & Murphy, 2020; Baker, 2016a, 2016b; Hemeda et al., 2014; Jung et al., 2010; Liu et al., 2023). Given the harsh conditions required to solubilize tumor matrix proteins, these hydrogels likely contain fragmented ECM proteins that are incapable of physiologic matrix self-assembly (Hughes et al., 2010; Tuftee et al., 2024).

Human cell–derived ECM represents a promising alternative, as cells can synthesize, process, and assemble matrix proteins into organized networks that more closely resemble native tissue architecture (Fonseca et al., 2023; Franco-Barraza et al., 2016; Muiznieks & Keeley, 2013; Solomonov et al., 2025). Approaches including tissue decellularization and cell-generated microtissues have demonstrated the feasibility of producing human ECM scaffolds *in vitro* (Chen et al., 2025; Chiang et al., 2021; Duisit et al., 2018; Fonseca et al., 2023; Jin et al., 2022; Johnson et al., 2016; Malakpour-Permlid et al., 2021; Zhang et al., 2023). Mesenchymal stromal cells (MSCs) are attractive candidates for matrix production due to their availability from multiple tissues, robust secretory capacity, and established use in biomanufacturing (Fonseca et al., 2023; Pompili et al., 2021; Smith et al., 2020; Tao et al., 2020). However, most studies of MSC-derived ECM have relied on animal-derived supplements such as fetal bovine serum (FBS) or human platelet lysate (hPL) (Amable et al., 2014; Chen et al., 2025; Maia et al., 2014; Novoseletskaya et al., 2020; Sagaradze et al., 2020; Yin et al., 2024).

The use of serum-based culture systems presents significant limitations for matrix manufacturing. Growth factors present in FBS and hPL become sequestered within the ECM, embedding xenogeneic or donor-dependent proteins that compromise matrix definition, reproducibility, and downstream applications (Baker, 2016a, 2016b; Hemeda et al., 2014; Jung et al., 2010; Keppie et al., 2021; Liu et al., 2023; Zhang et al., 2017). Chemically defined media offer a path toward reproducible, animal-component-free bioprocessing (Malakpour-Permlid et al., 2025; Oredsson et al., 2025; Weber et al., 2024), but these formulations lack the complex growth factor milieu found in serum and are often associated with reduced cellular performance (Hemeda et al., 2014; Jung et al., 2010; Spoerer et al., 2025; Xu et al., 2020). Critically, matrix production requires not only synthesis of ECM proteins, but also coordinated expression of processing enzymes necessary to convert precursor molecules, such as procollagens, into mature, self-assembling networks (Muiznieks & Keeley, 2013; Solomonov et al., 2025). Whether chemically defined media can support this level of MSC-mediated matrix assembly remains unknown.

Here, we investigate whether human MSCs from multiple tissue sources can self-assemble basement membrane–type matrices under fully defined, animal-component-free conditions. Using a quality-by-design approach compatible with GMP manufacturing, we show that MSCs cultured in chemically defined media form three-dimensional spheroids and adherent multilayered sheets that deposit Col IV–positive basement membrane matrices. These matrices contain canonical basement membrane components, can be decellularized, and support recellularization, demonstrating their potential as scalable and human-relevant ECM scaffolds for bioengineering applications.

## 2 Materials and Methods

### 2.1 Cell source

High quality human MSCs were obtained from commercial sources. UC were provided by Tissue Regeneration Therapeutics (TRT), Inc. (Toronto, Canada). UC MSCs were isolated from the perivascular Wharton’s jelly of healthy term (>37 weeks) umbilical cords from 1 male infant delivered by C-section and cryopreserved at mean population doubling level (mPDL) 4.87. AD and BM MSCs from female donors were obtained from RoosterBio Inc. (Frederick, MD) at 8.9 and 8.8 mPDL, respectively. PL MSCs from a female donor was obtained from OrganaBio (Miami, FL) at mPDL 8.39. Experiments were performed using one donor population per tissue of origin. For whole mount immunostained spheroids (Section 2.7), 3 UC MSC donor populations were pooled and cultivated.

### 2.2 Cell culture and maintenance

For cell expansion, MSCs were seeded at a density of 1,333 cells (UC MSCs) and 3,000 cells (AD, BM, PL MSCs)/cm^2^ in chemically defined phenol red-free MSC NutriStem XF^®^ complete growth media (Sartorius AG, USA). Cells were seeded according to TRT and RoosterBio’s recommended seeding density and expansion protocols which have been optimized for maximal cell expansion. Culture-ware was pretreated with 2.67µg/cm^2^ Col IV from human placenta (MilliporeSigma) to facilitate adherence. Cells were maintained in physioxia at 37°C, 5% CO2, 5% O2 and 80% relative humidity in a Heracell VIOS 160i tri-gas incubator (ThermoFisher Scientific, Waltham, MA). At 70-80% confluence, MSCs were washed briefly with Dulbecco’s phosphate buffered saline (DPBS; MilliporeSigma) and enzymatically detached by 2-3 minute incubation with TrypLE Select (ThermoFisher Scientific, Waltham, MA). Upon complete dissociation, an equal volume of media was added to the cell suspension and cells enumerated using the CellDrop Fluorescent Cell Counter (DeNovix Inc., USA). Cells were pelleted by centrifugation at 150 x *g* for 5 minutes and the cell pellet resuspended in fresh chemically defined media before re-seeding. All experiments were performed using MSCs between PDLs 16-24.

### 2.3 Self-assembled spheroids

Spheroids were generated using a protocol adapted from Li et al (Li et al., 2021): 96-well plates were coated with 75µl of sterile 1% agarose (w/v) and allowed to solidify at room temperature. Coated plates were used immediately or stored up to 1 week in 80µl DPBS per well. DPBS was aspirated prior to seeding. MSCs were suspended in NutriStem XF^®^ culture media at a concentration of 437,500 cells/ml (35,000 cells/well). 80µl of cell suspension was added to the coated wells and spheroids were allowed to spontaneously aggregate over 4 days in the incubator. To generate spheroids of different sizes, UC MSCs were cultivated using initial cell counts of 7,000, 15,000, 35,000, or 75,000 MSCs per well.

### 2.4 Morphometric analysis

To quantify differences in spheroid morphology across MSC tissue sources, AD, BM, PL and UC MSCs spheroids were nucleated using 35,000 cells/well. Phase contrast images were captured on days 1, 2 and 4 using an Echo Revolve inverted microscope (ECHO Inc., USA) with 10X objective for morphometric analysis. Images were acquired on days 1, 2 and 4 and processed using Adobe Photoshop 2023 (24.0.1, Adobe, USA). Spheroids were selected using Object Selection. In cases where the background was too high for auto-selection, the Magnetic Lasso tool was used to manually trace the spheroid perimeter. The background was replaced with grey as a separate layer using Photoshop to restrict calculations of each spheroid. For the initial spheroid seeding density experiments, the Ruler tool was used to measure spheroid diameter after 4 days in culture. Next, the Measure tool was used to automatically quantify spheroid area, perimeter and circularity. At least 19 spheroids per biological replicate, per tissue source were quantified.

### 2.5 Multilayered MSC sheets

MSCs were expanded and maintained to harvest an appropriate number of cells for experiments. High-density MSCs (7,813 cells/cm^2^) were seeded into 24 well plates in chemically defined media with media changes every 2-3 days and incubated at 37°C, 5% CO_2_, 5% O_2_ and 80% relative humidity. At specified timepoints (days 1, 2, 4 or 7 after seeding), MSCs were washed with DPBS and then decellularized (see section 2.6) for subsequent immunostaining.

### 2.6 Decellularization

Spheroids and multilayered sheets were washed twice in 1X DPBS for 5 minutes each then incubated in decellularization solution (20mM NH4OH + 0.5% Triton-X100 plus 1X cOmplete^TM^ EDTA-free protease inhibitor cocktail (Roche)) for 30 minutes at 37°C, 5% CO_2_, 5% O_2_ and 80% relative humidity. To remove residual DNA, the decellularized spheroids were washed twice more in DPBS, then treated with 50kU of DNAse I plus EDTA-free protease inhibitor diluted in water for 30 minutes at 37°C, 5% CO_2_, 5% O_2_ and 80% relative humidity. The washes were repeated and then the spheroids were stored at 4°C for whole mount immunofluorescence or submerged in 75µl radioimmunoprecipitation (RIPA) buffer and immediately flash frozen in liquid nitrogen and stored at -80°C until use for western blot analysis.

To assess retention of cellular material after decellularization, untreated MSC sheets were immunostained with anti-carboxymethyl lysine (anti-CML; 1:200; ab125145; Abcam, USA) as detailed in Section 2.7. The sheets were imaged, then decellularized and re-imaged (Fig S1A). All images were captured using a Nikon ECLIPSE Ti2-E inverted microscope with the 20X objective lens.

Retention of nuclear material after decellularization was assessed using FIJI analysis of DAPI-stained decellularized matrices (Fig S1.B). DAPI 16-bit images were imported and threshold with black background was set for the cellularized (913-65535) and decellularized (1600 or 2724-65535) respectively. A gaussian blur with sigma factor 2 was applied with auto thresholding set to “default dark”. The images were masked based on the thresholding values, and the “analyze particles” tool was used to obtain nuclei count. Intact nuclei were counted from at least 10 fields of view for each of 3 decellularized wells and 1 cellularized well in each experiment (Fig. S1C). Decellularization efficiency was calculated as the percent of nuclei in decellularized wells compared to cellularized wells.

### 2.7 Immunofluorescence

Spheroids and sheets were immunostained in 1.5ml microcentrifuge tubes with nutation or in-well, respectively. For confocal imaging, UC MSCs were seeded at 4000 cells/cm^2^ onto coated IBIDI 8-well chamber slides. Cellularized samples were fixed in 10% neutral buffered formalin (NBF; MilliporeSigma) for 10 minutes, rinsed in 1X DPBS and then permeabilized with PBT (1X DPBS + 0.1% TritonX-100) for 10 minutes. Following 3 washes in 1X DPBS, samples were blocked with 10% normal donkey serum (MilliporeSigma) diluted in PBST (PBS + 0.1% Tween-20) for 30 minutes. Primary antibodies against human collagen IV (Col IV, 1:200, NovusBio NB120-6586; Col IV alpha 1 NC1, 1:100 Chondrex 7070), laminin (LM, 1:200, NovusBio NB300-144), fibronectin (FN, 1:200, NovusBio AF1918) collagen VI alpha 1 (Col VI, 1:200, NovusBio NB120-6588) or lysl oxidase-like 2 (LOXL2, 1:100, Abnova 675-773) were diluted in antibody dilution buffer (PBS + 0.1% Triton-X100 with 1% bovine serum albumin (BSA)) (MilliporeSigma) and incubated with the sample overnight at 4°C. Scaffolds were washed 3 times for 5 minutes each with 1X DPBS at room temperature, then incubated for 1 hour at room temperature with secondary antibodies (donkey anti-Rabbit IgG (H+L) conjugated to AlexaFluor^TM^ 647 (1:1000, Invitrogen A32795), (goat anti-Rat 1gG (H+L) conjugated to Dylight^TM^ 680 (1:500, Novus NBP2-68483FR)) or donkey anti-Sheep IgG conjugated to AlexaFluor^TM^ 594 (1:1000,Invitrogen A110106)) in antibody dilution buffer. The samples were rinsed twice with 1X DPBS and then stained with 4’,6-diamidino-2-phenylindole (DAPI, 1:4000, MilliporeSigma) for 2 minutes at room temperature, followed by three 5 minutes each in 1X DPBS. Spheroid samples were stored in Invitrogen Cytovista 3D tissue clearing reagent (ThermoFisher Scientific, Waltham, MA) until mounting. These methods necessitated the use of common animal-derived antibodies and blocking solutions; where possible, we are working to use alternatives like human serum albumin going forward.

Whole mount imaging of spheroids was facilitated using platform-type slides to accommodate the sample depth and prevent deformation under coverslip pressure. No.2 coverslips were adhered to either end of the microscope slide using clear nail polish to create elevated platforms. The immunostained samples, suspended in Cytovista solution, were pipetted onto the slide and overlaid with a 22x40mm No. 1 coverslip. Mounted slides were sealed with clear nail polish and allowed to set overnight before imaging. Confocal images were captured using a Nikon A1R confocal microscope equipped with PlanFluor 10X DIC L N1 and 40X Oil DIC H N2 lenses. Images were analyzed using NIS Elements software (Nikon Instruments, Inc., USA).

Epifluorescent images of matrix deposition in adherent sheets were captured using a Nikon ECLIPSE Ti2-E inverted microscope with the 20X objective lens with 32X averaging on NIS Elements software (Nikon Instruments, Inc., USA). Images were edited using FIJI (v1.54k; (Schindelin et al., 2012)). Wells corresponding to days 1 and 2 were edited to remove background and visualize weak signal. Raw images were imported into FIJI as 16-bit, greyscale images and the brightness and contrast of each image optimized: FN minimum and maximum levels were set to 185 and 400, respectively; Col IV minimum and maximum levels were set to 170 and 600, respectively; Col VI minimum and maximum levels were set to 120 and 275, respectively; LM minimum and maximum levels were set to 115 and 540, respectively. Images taken on days 4 and 7 were comparably adjusted to maintain consistency between timepoints. FN minimum and maximum levels were set to 160 and 250, respectively; Col IV minimum and maximum levels were set to 145 and 275, respectively; Col VI minimum and maximum levels were set to 125 and 200, respectively; LM minimum and maximum levels were set to 115 and 190, respectively.

### 2.8 Western blot

Total protein was collected from at least 15 pooled, decellularized and DNAse I-treated spheroids per tissue source and replicate. 3D matrices were solubilized by 3 rounds of sonication at 10% amplitude for 30 seconds (1 second on, 1 off). Protein concentration was determined using Bicinchoninic acid assay (BCA; MilliporeSigma) for protein normalization. Samples were diluted and prepared in 6X Laemmli buffer to load 3ug of total protein in 10µl per lane then boiled at 95°C before loading on an 8% gel (or 12% for elastin). Proteins were subjected to electrophoresis for 1.5 hours at 140V. Size fractionated proteins were transferred to a nitrocellulose membrane overnight at 4°C at 30V. Membranes were washed three times in 1X Tris Buffered Saline (TBS) for 10 minutes each and then blocked for 1 hour at room temperature in 5% skim milk, or 5% BSA for elastin. The membranes were then incubated with primary antibody solution (rabbit anti-human col IV (1:1000, NovusBio, NB120-6586), rabbit anti-laminin (1:1000, NovusBio, NB300-144), rabbit anti-col VI alpha 1 (1:2000, NovusBio NB120-6588), sheep anti-fibronectin (1:3000, NovusBio AF1918) or rabbit anti-elastin (1:3000, Invitrogen PA5-99418) overnight at 4°C.

Membranes were then washed three times in 1X TBST for 10 minutes each. Secondary antibodies (goat anti-rabbit IgG, H+L chain specific peroxidase conjugate (1:3000, Millipore Sigma 401393), rabbit anti-sheep IgG (H+L) secondary antibody (1:4000, Invitrogen 61-8620) or AMDEX™ goat anti-rabbit horseradish peroxidase conjugate (1:3000, Cytiva RPN4301) were diluted in blocking solution and added to the membrane for 1 hour at room temperature. Membranes were washed four times in 1X TBST and visualized using UltraScence Pico Ultra Western Substrate (FroggaBio). Western blots were imaged using an Amersham^TM^ AI600 Imager (GE HealthCare Technologies, USA).

### 2.9 Recellularization of matrix sheets

Recellularization of decellularized matrix sheets was assessed using a sample of pooled UC MSCs from 3 donors. Cultured MSCs in T25 tissue culture flasks were pre-stained with Calcein-Green AM (1:1000; ThermoFisher Scientific, Waltham, MA) for 10 minutes, rinsed with 1X Hanks’ Balanced Salt Solution with Ca^2+^ and Mg^2+^, dissociated using 1X TrypLE and centrifuged as described in Section 2.2. Calcein-Green labeled MSCs were resuspended at a concentration of approximately 3900 cells/ml in culture media. Unfixed MSC-derived sheet matrices were pre-labeled using anti-FN antibody as described in Section 2.7. Cell suspension was added at a volume of 500 μl/well for 24 well plates and 250 μl/well for 48 well plates and incubated overnight as described in Section 2.2. The next day, the culture vessels were placed in a temperature- and humidity-controlled stage top incubator (TOKAI HIT, Co., Ltd., Japan). Time lapse images of cell-matrix interactions were captured every 5 seconds over a 2-minute period using the Nikon ECLIPSE Ti2-E inverted microscope (Nikon Instruments, Inc., USA) under a 20X objective.

### 2.10 Statistical analyses

Statistical analysis was carried out using Prism 10.3.1 (GraphPad Software, LLC). Significant differences were calculated using student’s t-test (decellularization efficiency), 2-way analysis of variance (spheroid morphometrics) and Kruskal-Wallis test (spheroid initial seeding density). Error bars represent standard deviation (SD) ± of the calculated means. ns = non-significant, P>0.05, * P ≤ 0.05, ** P ≤ 0.01, *** P ≤ 0.001 and **** P ≤ 0.0001. All experiments were performed 3 independent times, exception immunostaining of whole mount spheroid scaffolds which was performed using 2-3 technical replicates.

## 3 Results

### 3.1 MSCs from different human tissues self-assemble into 3D spheroids in chemically defined media

We first asked whether MSCs from adult (AD, BM) and perinatal (PL, UC) tissues could reliably form spheroids with an organized matrix in animal component-free, chemically defined conditions. Within four days of high-density seeding on agarose-coated 96-well plates, MSCs from both adult and perinatal tissues formed a single free-floating spheroid in each well (Fig. 1A). A small number of unincorporated cells remained in solution and formed micro-aggregates or attached to the wall of the well, particularly in AD MSC cultures (not shown).

**Figure 1:**
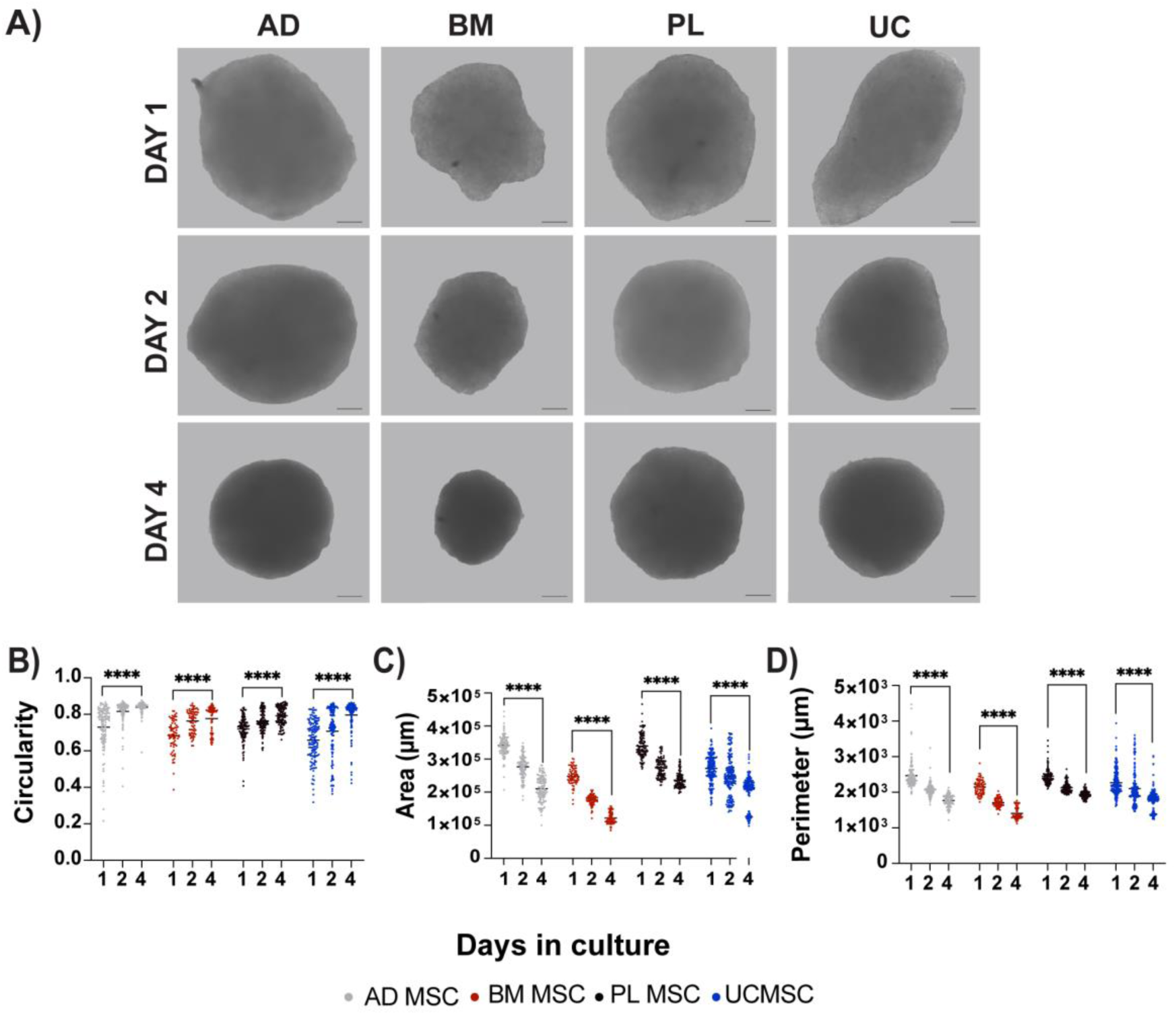
Scaffold-free MSC spheroids exhibit features of self-directed organization. Morphometric analysis of self-assembled MSC spheroids in defined media reveals (A) a visible increase in uniformity which can be quantified by (B) increased circularity (1.0 is a perfect circle), (C) reduced area and (D) decreased perimeter, which together are hallmarks of tissue compaction. Scale bar = 100µm. **** p< 0.0001.

Spheroid geometry influences ECM synthesis and organization both directly (Gonzalez-Fernandez et al., 2022) and indirectly by signaling pathways associated with nutrient and oxygen transport (Kumar et al., 2018; Murphy et al., 2017; Shearier et al., 2016), cell–cell interactions (Im et al., 2021) and cytoskeletal tension (Zhou et al., 2017). Accordingly, we quantified spheroid morphometrics (diameter, area, and circularity) over time to assess whether changes in matrix content are associated with, or potentially driven by, alterations in spheroid structure.

Over time, the spheroids became increasingly uniform in size and morphology. Circularity (where 1.0 is a perfect circle) increased from Day 1 to Day 4 by 11-15% for each MSC type, resulting in average circularities of 0.84, 0.78, 0.80 and 0.80 by Day 4 for AD, BM, PL and UC MSCs, respectively (Fig. 1B). Circularity was accompanied by compaction, evidenced by a significant decrease in area (Fig. 1C) and perimeter (Fig. 1D). Spheroids that were much smaller or larger than the mean on Day 1, which may not have aggregated as efficiently or in which there were cell dispensing errors, remained outliers over the time course although their morphometric changes followed the other spheroids.

Thus, we next investigated how initial cell density might influence spheroid self-assembly. Spheroids were nucleated using 7,000, 15,000, 35,000, and 75,000 MSCs per well (Fig. 2A). Irrespective of seeding density, all spheroids successfully self-aggregated. Spheroid size increased relative to seeding density, but not linearly (Fig. 2B). All spheroids were visible by eye with diameters ranging from 326 to 707µm (Fig. 2A). Excluding the largest aggregates nucleated with 75,000 MSCs, spheroids could be aspirated using a P1000 pipette without shearing. On the other hand, the smallest spheroids were challenging to visualize and manipulate. Considering these practical limitations, as well as reported oxygen diffusion limits (Murphy et al., 2017), we selected spheroids nucleated using 35,000 MSCs and cultured to Day 4 for the remainder of the study.

**Figure 2:**
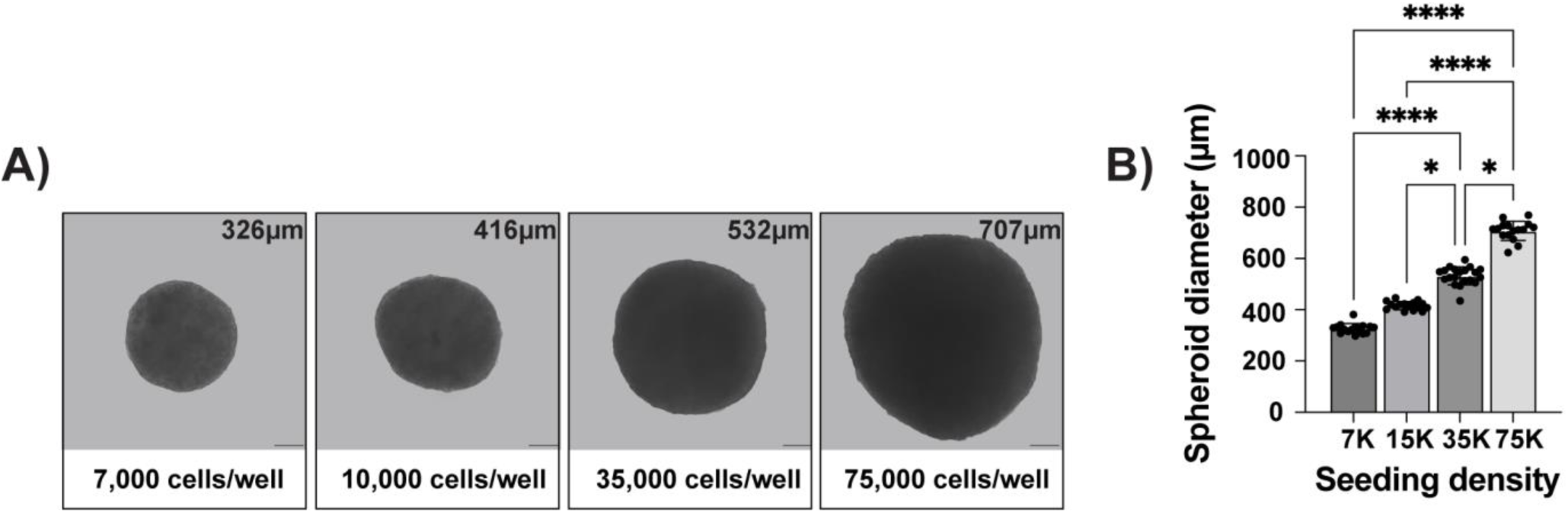
Spheroid diameter correlates with seeding density. Spheroids were seeded in defined media using 7,000, 15,000, 35,000 or 75,000 UC MSCs and measured after 4 days. (A, B) Spheroid size correlated with seeding density, but not in a linear fashion. Average spheroid diameter (μm) (A, upper right corner) was calculated from (B) 15 spheroids per group. Spheroid size was highly consistent between replicates. Representative images are from n=3 independent experiments. Scale bar = 100μm. * p< 0.5, **** p< 0.0001.

### 3.2 MSC spheroids produce Col IV-positive matrices in chemically defined media

We then asked whether the MSC spheroids cultivated in defined media contained basement membrane-type Col IV networks. Spheroids provide a physiologically relevant culture context in which 3D cell-cell interactions and biomechanical cues more closely resemble the tissue environment, in contrast to rigid 2D substrates that can bias MSCs towards fibrotic matrix production (Loomis et al., 2022; Park et al., 2023; Walker et al., 2021). Moreover, the spheroid format enables assessment of whether MSC-secreted ECM proteins are appropriately processed and assembled into organized, tissue-like basement membrane scaffolds.

Spheroids were decellularized using a protocol reported to preserve matrix architecture (Li et al., 2021) and immunostained for fibronectin (FN) and Col IV. Multi-labelling for additional basement membrane constituents was impeded by the lack of reliable commercial antibodies produced in other species. Confocal imaging of the spheroids at low magnification (10X), confirmed the tissue-dependent size differences documented by light microscopy (Fig. 1), but revealed complex surface topology, some of which may be an artefact of deformation after mounting which implies viscoelasticity (Fig. 3A). At higher magnification (40X), complementary networks of fibronectin and Col IV were observed (Fig. 3B). The decellularized matrices all presented architectural basement membrane features including pores, fibre alignment and rough surface topography (Fig. 3A), which are more clearly depicted in digitally enlarged representative images (Fig. 3B).

**Figure 3.**
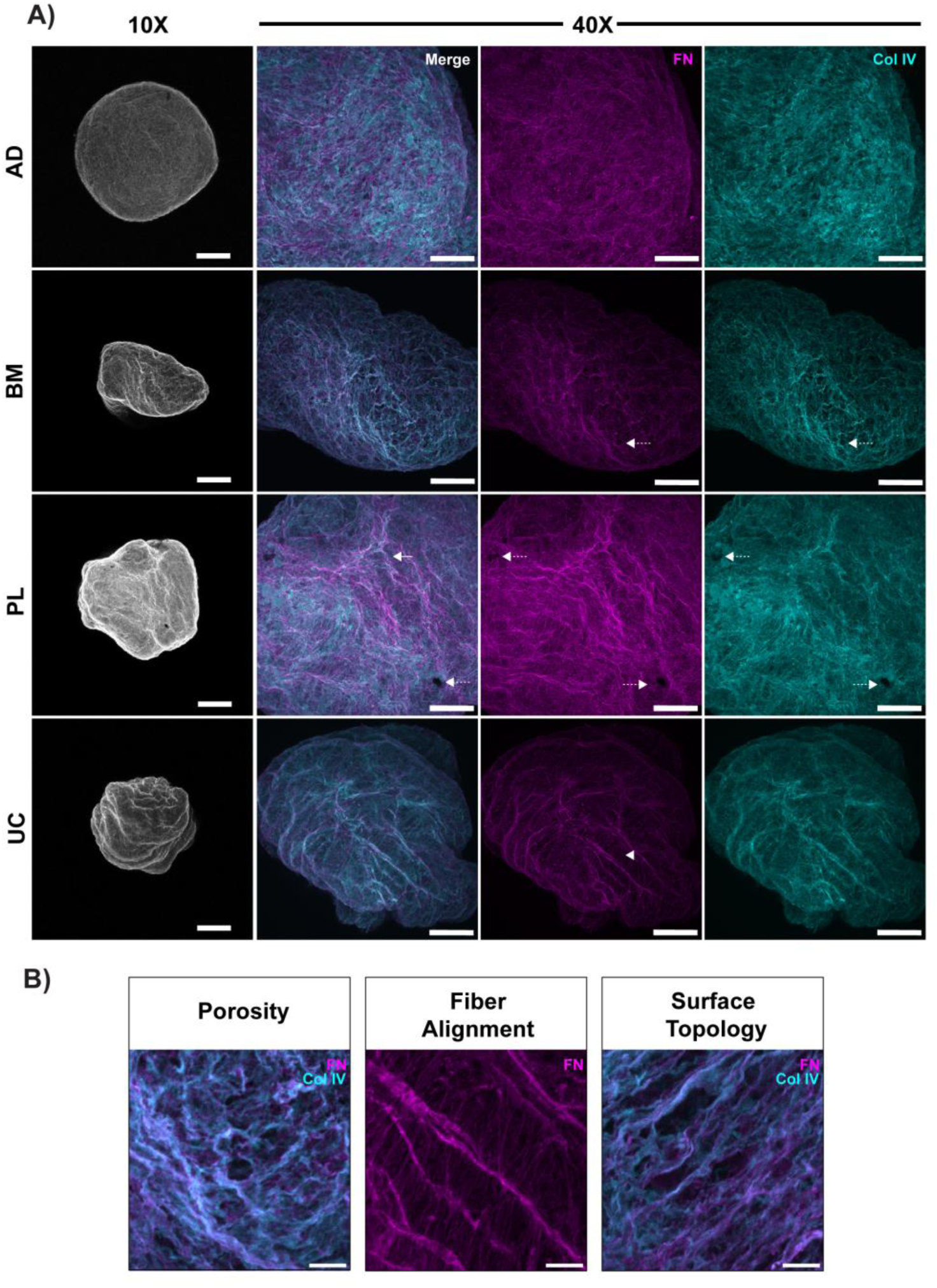
MSC spheroids produce Col IV-positive matrices in chemically defined media. Whole mount confocal images of 4 day old decellularized matrices produced by spheroid cultures of MSCs from adult (AD, BM) and perinatal (PL, UC) tissues. Matrices were immunostained for fibronectin (FN, magenta) and Col IV (cyan). (A) Maximum intensity projection of 2μm confocal images captured over a Z depth of 50μm with a 10x objective. Overlaid FN and Col IV confocal images, shown in greyscale, reveal surface roughness. Some loss of circularity and folding may be due to compression between the slide and the coverslip, suggesting that the scaffolds have plasticity. Scale bar = 50μm. Maximum intensity projection of 0.5μm slices captured over a Z depth of 80μm with a 40x objective reveal complex networks of FN and Col IV. These proteins appear to be aligned in a complementary fashion that moderately overlaps (white). Architectural features including pores (dotted arrow), fiber directionality (arrowhead) and surface topology (arrow) are also apparent. (B) Digital magnification of representative panels in (A) show enhanced architectural details including defined pore structures (Porosity), alignment of fibronectin fibers across 2 planes (Fiber Alignment), and rough Surface Topography featuring peaks and troughs. These topological and topographical features are found in all examined spheroid-derived matrices. Representative images from 3-4 technical replicates per MSC tissue source. Scale bar = 100μm.

Three-dimensional volumetric reconstruction of the confocal images revealed distinct patterns of matrix organization. In the adult MSC-derived matrices, fibronectin and Col IV appeared to be interwoven, while the perinatal MSC-derived matrices segregated fibronectin into apical sheets, with an underlying sheet of Col IV (Fig. 4A). Regardless of their relative orientation, fibronectin and Col IV rarely overlapped except at the interface between the apical and luminal matrix facets (Fig. 4B).

**Figure 4.**
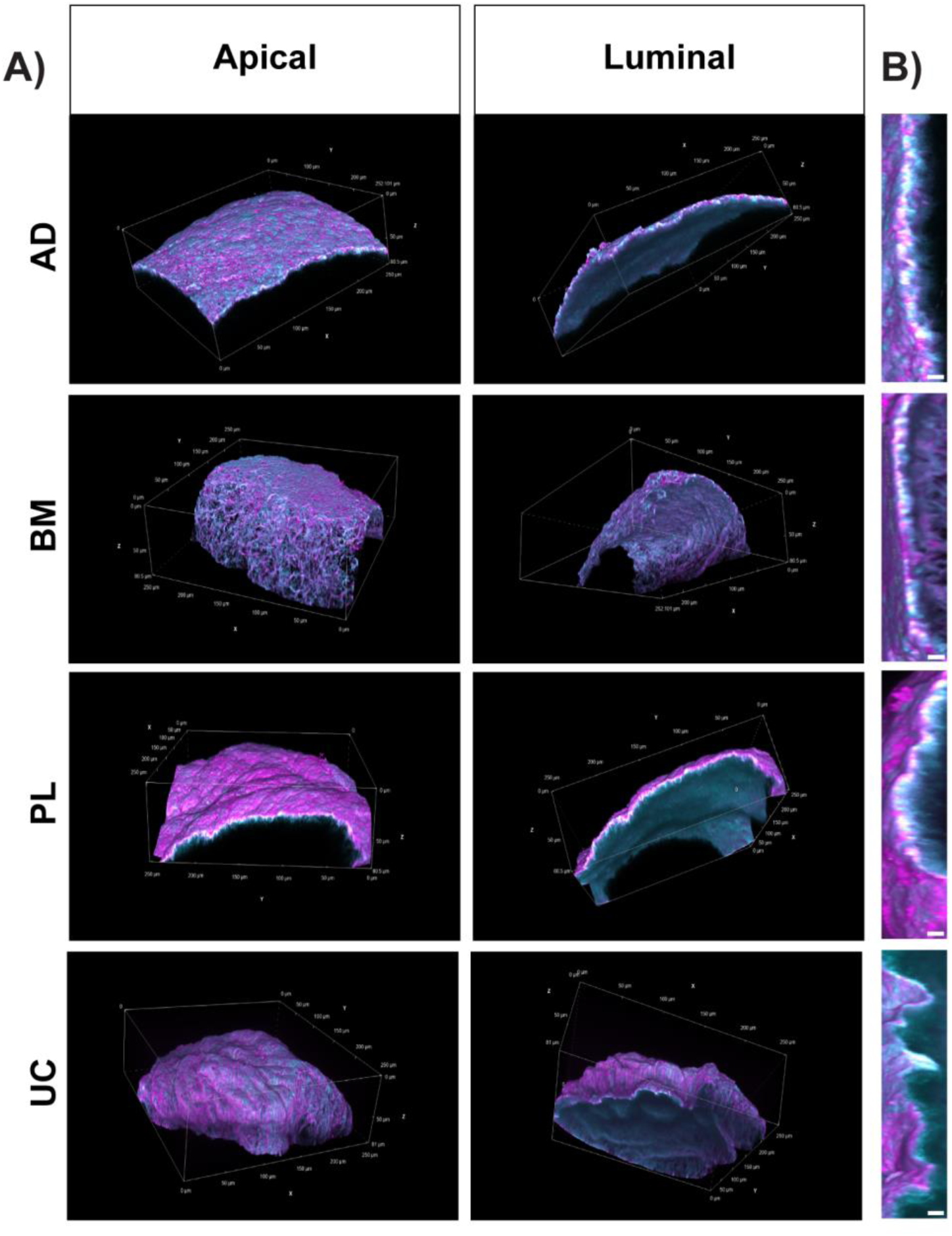
Perinatal-derived MSCs produce self-assembled matrices with overlaid FN and Col IV sheets. Decellularized whole mount spheroid-derived matrices produced by AD, BM, PL and UC MSCs were examined in volume view with alpha blending to highlight surface and compositional topography. (A) Fibronectin (FN, magenta) and Col IV (cyan) appear to be interwoven in matrices produced by adult MSCs (AD, BM). By contrast, the perinatal MSCs (PL, UC) establish matrices with sheet-like organization, whereby FN is enriched on the apical surface, complemented by a sheet of Col IV on the underlying luminal surface. (B) Cross-sections reveal a marked overlap of FN and Col IV (white) at the interface of the apical and luminal sheets. Scale bar = 5μm. Representative images from 3-4 technical replicates per MSC source. Confocal images were captured in 0.5μm sections through a Z-depth of 80μm.

### 3.3 Self-assembled MSC matrices contain canonical basement membrane factors

Having shown that MSC-derived matrices contain networked fibronectin and Col IV, we used western blot to assess whether other basement membrane factors were also present. Decellularized spheroids were solubilized, normalized to total protein, size fractionated and then immunoblotted for fibronectin, laminin β/γ, Col IV, Col VIα1 and elastin (Fig. 5). Laminin β and γ chains, along with an α chain, form the basic heterotrimeric structure of all laminins. They assemble with other proteins, such as Col IV and nidogen, to create the complex, organized network of the basement membrane (Lennon & Sherwood, 2025; Page-McCaw & Ferrell, 2025; Page-McCaw et al., 2025). Col VIα1 is a key component of type VI collagen, which acts as a structural link between the basement membrane and the surrounding interstitial matrix, providing mechanical support to tissues (Cescon et al., 2015; Gregory et al., 2024). Elastin anchors cells to the matrix and provides mechanical support; it functions in conjunction with other basement membrane components to provide elasticity, allowing tissues to stretch and recoil (Krymchenko et al., 2025; Ruiz et al., 2024; Trębacz & Barzycka, 2023). The relative abundance of these analytes differed in matrices produced by each MSC donor. BM MSC-derived matrices were markedly enriched for fibronectin, while UC MSC-derived matrices were enriched for all the basement membrane constituents compared to the other donors (Fig. 5). Col IV and elastin were less abundant in the AD-derived scaffolds, while all the analytes except elastin were detected in PL-derived matrices (Fig. 5).

**Figure 5:**
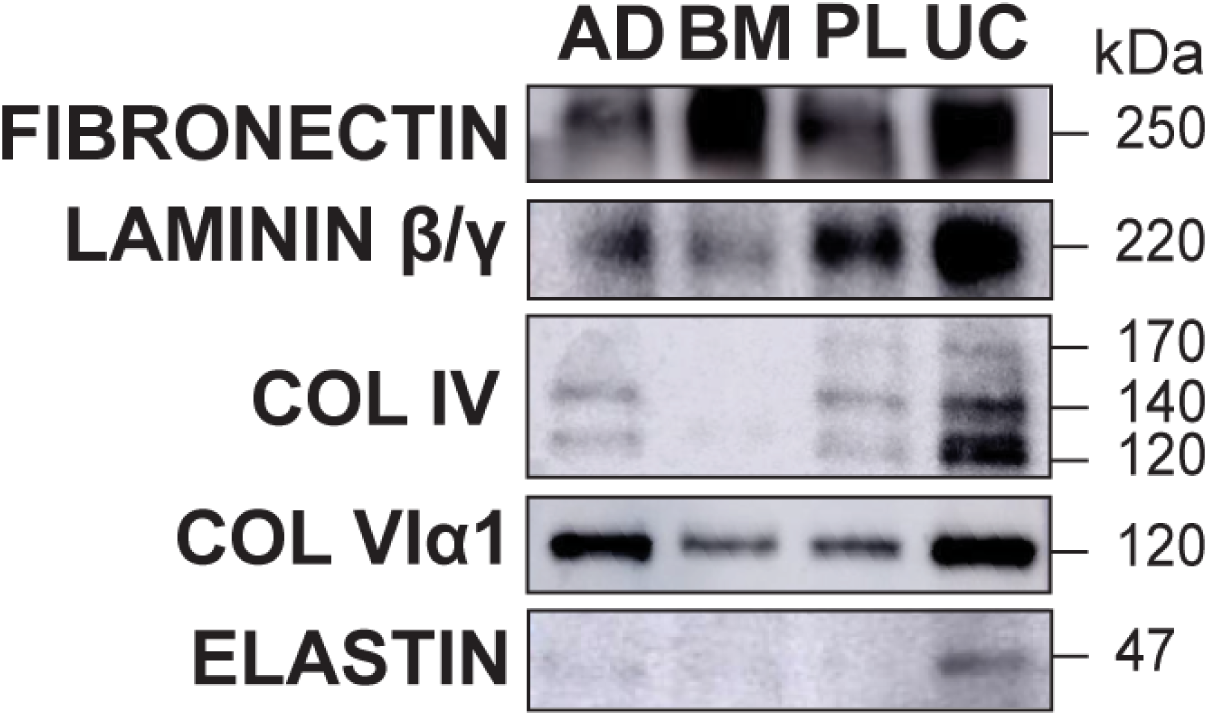
MSC-derived matrices contain basement membrane-specific proteins. Relative abundance of basement membrane proteins in decellularized AD, BM, PL or UC MSC spheroids were analyzed by western blot. Samples were normalized by total protein (3μg/lane). UC MSC-derived matrices were enriched for the basement membrane factors Col IV, Col VIα1 and elastin. Matrix from BM MSCs contained relatively more fibronectin than the other scaffolds. Representative data from n=3 independent experiments.

### 3.4 Multilayered UC MSCs produce matrix basement membrane-type sheets in defined media

Having determined that the UC MSC donor produces synthesizes the most robust repertoire of basement membrane proteins when cultured as spheroids (Fig. 5), we next asked whether a similar basement membrane-type matrix could be generated when UC MSCs were grown as adherent cultures. Compared with spheroids, multilayered cell sheets are technically simpler and may offer greater reproducibility and scalability. However, adherence to rigid culture substrates imposes a stiffer microenvironment and reduces 3D cell-cell interactions, conditions that are known to influence matrix deposition and may preferentially promote interstitial rather than basement membrane matrix production. We therefore sought to determine whether basement membrane assembly is preserved under these 2D culture conditions.

UC MSCs were cultured in adherent multilayers followed by decellularization to assess ECM biogenesis. We then analyzed the deposition of select adhesive (fibronectin, laminin β/γ) and structural (Col IV, Col VIα1) basement membrane proteins over time by immunofluorescence. Fibronectin was the most prevalent matrix component at Day 1 and increased in abundance and architectural complexity through Day 7 (Fig. 6A, B). Col IV and Col VIα1 were barely detectable at Day 2 and continued to accumulate in networked structures through Day 7 (Fig. 6A, B). At Day 7, laminin β/γ was the last analyte to be detected. The matrix sheets were 8-10µm thick on Day 7, as determined using cross-sections of Z-stacked images (Fig. 6C). In cellularized samples, networked fibronectin (red) is seen connecting with and intercalating between MSCs (white) highly expressing the Col IV cross-linking enzyme lysyl oxidase-like 2 (LOXL2; Fig. 6C, D).

**Figure 6:**
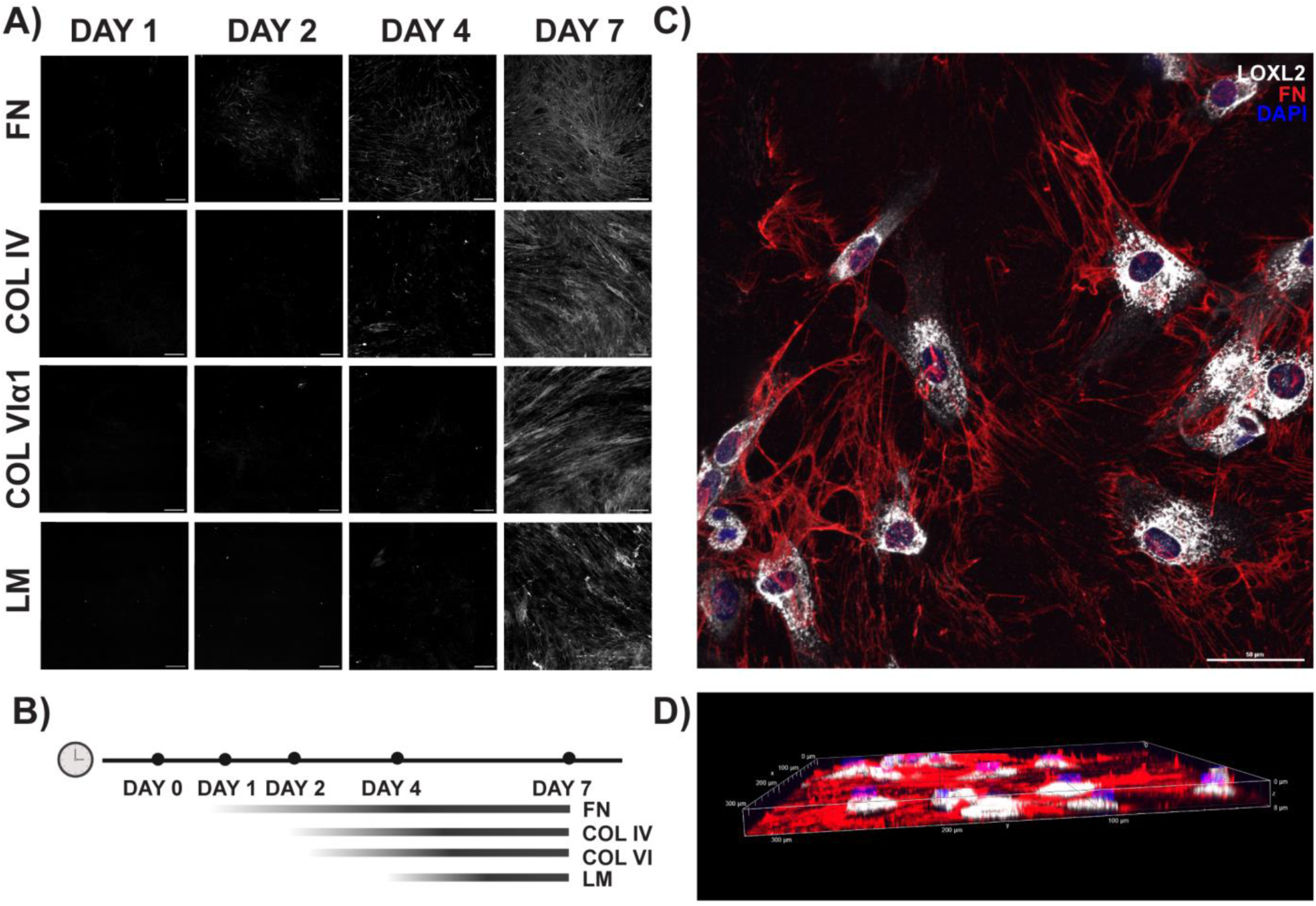
Multilayered UC MSCs construct sheets of scaffolded matrix. (A) Fibronectin (FN) is first detected by immunogenic staining 2 days after seeding, followed by Col IV on Day 4. Col VIα1 and laminin (LM) are not readily detected until Day 7, when all 4 analytes are visible components of a networked matrix structure. Scale bar = 100μm (B) Schematic summarizing the temporal incorporation of ECM components depicted in (A). (C) Maximum intensity projection 4-day MSC multilayers show networks of fibronectin (FN; red) intercalating between MSCs expressing high levels of the Col IV cross-linking enzyme LOXL2 (white), in (D) a matrix 8μm thick. Data shown is representative from a dataset of n=3 independent experiments with 5-6 fields of view per experiment.

### 3.5 UC MSCs matrix sheets are amenable to recellularization

Finally, we asked whether MSC-derived matrices were amenable to recellularization and could support cell-matrix interactions. Matrix sheets proved more practical and efficient than spheroids for reseeding and were selected for this experiment. Fluorescent labeled UC MSCs were incubated overnight with decellularized matrix sheets immunostained to visualize fibronectin. The next day, cell-matrix interactions were assessed by time lapse imaging. MSCs successfully adhered to the nascent scaffolds (Fig. 7). Time-lapse microscopy showed MSCs attached to the fibronectin fibers. MSCs appeared to leverage this matrix interaction to migrate through focal planes of the matrix (Fig. 7A) by pulling or tugging on the flexible fibronectin fibers (Fig. 7B, Movie S1). In preliminary studies, matrix produced by MSC in our animal component-free workflow can support the adhesion, migration and proliferation of other cell types including human neural progenitor cells, human umbilical vein endothelial cells and fibroblasts (not shown). Taken together, this study establishes MSCs as a viable and ethically favorable source for the generation of self-assembling matrix proteins, including basement membrane components, in chemically defined conditions.

**Figure 7:**
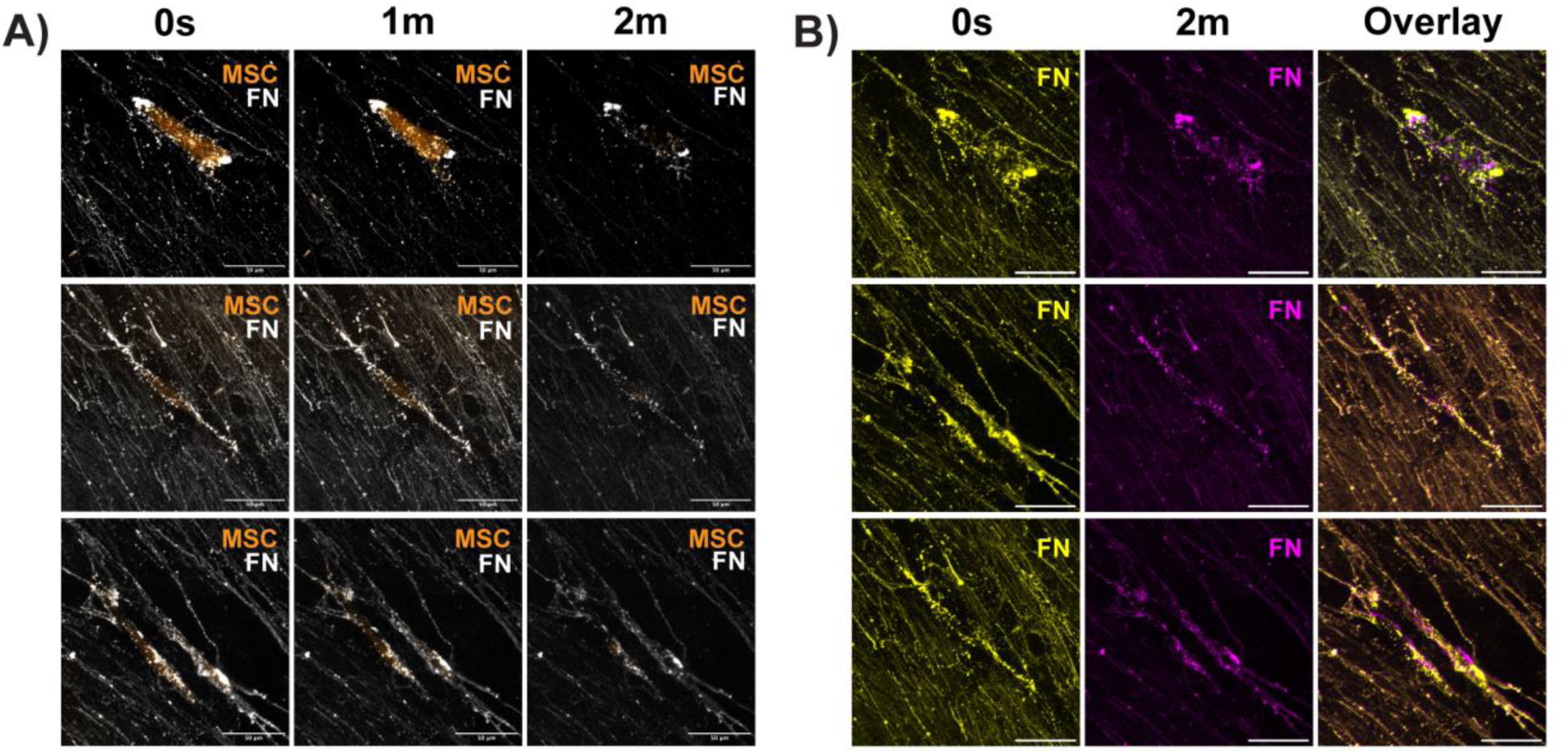
UC MSC derived matrix sheets can be colonized by new cells. Decellularized UC MSC sheets, produced in chemically defined media, support interactive cell migration of newly introduced UC MSCs **(Movie S1).** (A) In representative time lapse images, MSCs (orange) can be seen moving through the Z plane of scaffolded fibronectin (FN; white). FN scaffolds were false colored at 2 timepoints (0s, yellow; 2m, magenta) and overlaid to detect FN movement. The presence of unblended FN signal in cell-occupied regions shows relocation of FN, indicating dynamic cell-matrix interactions. Blended color (orange) in the unoccupied scaffold areas suggests that FN remains static in the absence of cell movement. Scale bar = 50µm.

**Figure S1:**
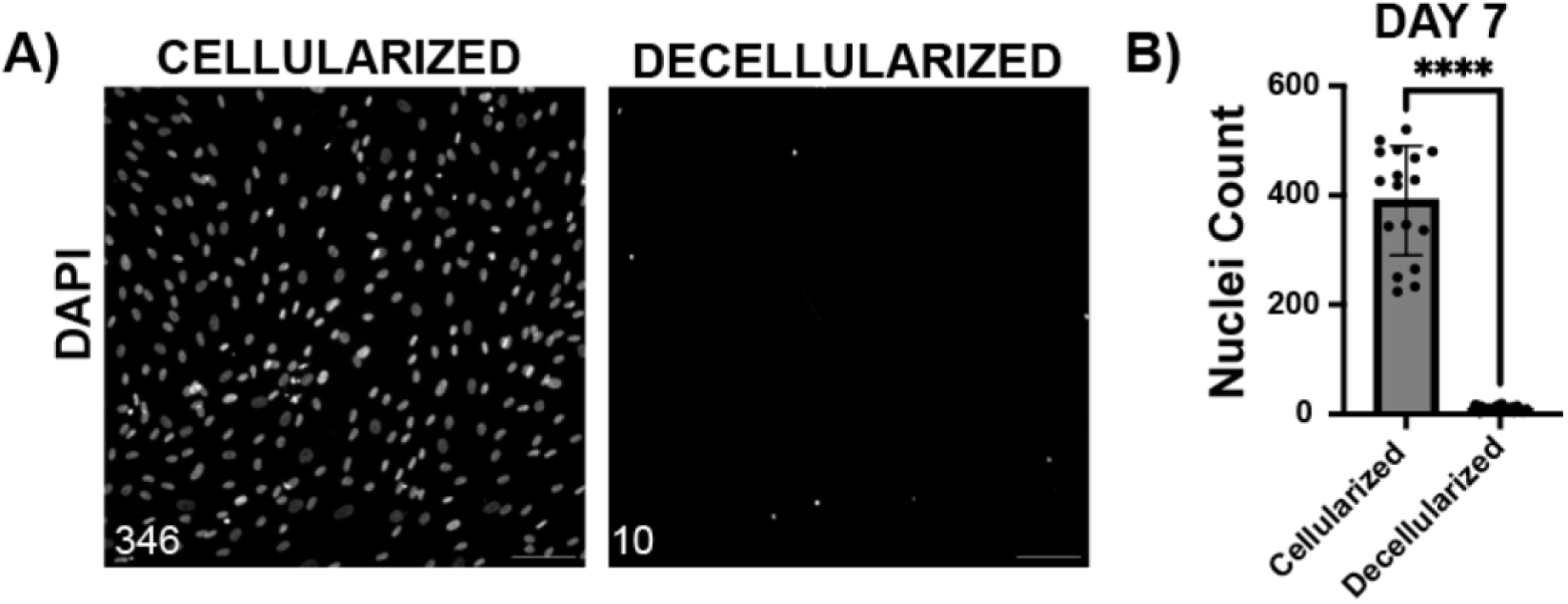
Ammonium hydroxide decellularization efficiently clears nuclear and cytoplasmic material. Multilayered UC MSC sheets were decellularized on Day 4 (A) and Day 7 (C) after seeding using an ammonium hydroxide protocol. A) Adhered UC MSCs were labeled using DAPI (nucleus) and anti-CML antibody to detect <u>c</u>arboxy<u>m</u>ethyl <u>l</u>ysine residues in the nucleus and cytoplasm. Negligible signal was detected on either channel following ammonium hydroxide-based decellularization, indicating efficient clearance of nuclear and cytoplasmic material. (B) Nuclei were labeled using DAPI and quantified (number in the bottom left of the image). (C) The number of nuclei was significantly reduced by 95.8% after decellularization. Representative data from n=3 independent experiments. Scale bar = 100µm.

## 4 Discussion

There is a critical need for functional human matrix components to support the next generation of high-fidelity tissue models for experimental use and transplant (Catarino et al., 2022; Liu et al., 2023; Naderi-Meshkin et al., 2023; Schneider et al., 2023). Moreover, there is a paucity of intact and functional basement membrane biomaterials to recapitulate the complexity of human tissues, and more importantly, to facilitate vascularization (Chan et al., 2016; Marrella et al., 2018; Minor & Coulombe, 2020; Zhao et al., 2022). Here we report that human MSCs, which occupy perivascular niches *in vivo* (Caplan, 2017; Corselli et al., 2013; Kang et al., 2010), can be leveraged to synthesize functional basement membrane components in a defined, animal component-free bioprocess. The development of an animal-component-free platform based on human MSCs offers a renewable and physiologically relevant strategy for scalable ECM production, reducing reliance on ethically contentious animal-derived materials, lowering the environmental burden associated with animal agriculture, and minimizing the risk of xenogeneic contamination.

Our findings align with previous studies that demonstrate origin-specific synthesis of ECM factors by MSCs, although this study is not sufficiently powered to statistically correlate MSC tissue source with ECM composition. In previous transcriptome studies that span multiple MSC donors and origins, we found that UC MSCs express significantly higher levels of the non-fibrillar basement membrane collagens (Col IV and Col VI) than fibroblasts (Wiese & Braid, 2020) and BM MSCs (Wiese et al., 2022). Similarly, Ragelle et al., 2017 found that decellularized neonatal dermal fibroblast-derived matrices possessed less total collagen than AD and BM MSC-derived scaffolds. The most abundant structural components in those BM MSC-derived scaffolds were Col I and Col VI, while Col II, III and IV were present at lower levels (Ragelle et al., 2017). These results were consistent across all 5 donors (Ragelle et al., 2017). Another group reported that decellularized BM MSC scaffolds contained Col IV, while AD MSC scaffolds did not (Marinkovic et al., 2020), which contradicts our findings. Amable *et al*. compared ECM components secreted by AD, BM and UC MSCs, finding that UC MSCs did not produce detectable amounts of laminin, fibronectin, Col II, or IV. In their study AD MSCs were the only MSC population to secrete Col IV at detectable concentrations, and they also produced more Col I, II and III than BM and UC MSCs (Amable et al., 2014). It is plausible to expect that these disparate outcomes are the consequence of different media formulations used in these studies, including FBS supplemented, (Amable et al., 2014; Bartosh et al., 2010; Cesarz & Tamama, 2016; Chen et al., 2025; Chiang et al., 2021; Fuentes et al., 2022; Lee et al., 2016; Marinkovic et al., 2020; Ragelle et al., 2017; Yin et al., 2024), serum-free (Choi et al., 2021; Rakian et al., 2015), and xeno-free products (Bauman et al., 2018; Parvin Nejad et al., 2023). This study aimed to test whether MSCs are capable of synthesizing basement membrane–type matrices under chemically defined culture conditions that exclude the rich array of growth factors, adhesion factors, and stabilizing components found in serum or hPL-supplemented media (Delabie et al., 2025; Hodges & Melcher, 1976; Karnieli et al., 2017; Klinker et al., 2017; Oikonomopoulos et al., 2015).

In mammals, Col IV is encoded by a family of six genes (*COL4A*1–6) that produce discrete Col IV α-chains (α1–α6) (Page-McCaw & Ferrell, 2025; Page-McCaw et al., 2025). These α-chains can give rise to three supramolecular scaffolds: Col IV^α121^, Col IV^α345^, and Col IV^α121–α556^, which form networks with different structural and biomechanical properties (Page-McCaw & Ferrell, 2025; Page-McCaw et al., 2025). Expression of the *COL4A* genes differs between tissues and shifts during development, resulting in the formation of functionally specialized basement membranes throughout the body (Page-McCaw & Ferrell, 2025; Page-McCaw et al., 2025). The observation that MSCs from different tissues produce distinct Col IV α-chains was therefore not unexpected and is consistent with published reports. For example, we previously compared the transcriptome profiles of 3 donors each of UC and BM MSCs and their responses to cytokine stimulation. In that study, unstimulated BM MSCs, as used here, expressed very high levels of *COL4*A1 and *COL4*A2 and very low levels of *COL4*A5, and did not express *COL4*A3, *COL4*A4 or *COL4*A6 (Wiese et al., 2022). UC MSCs expressed even higher levels of *COL4*A1 and *COL4*A2, moderate levels of *COL4*A5 and low levels of *COL4*A3, *COL4*A4 and *COL4*A6 (Wiese et al., 2022). Notably, expression of these genes did not vary significantly between donor populations and were consistent with an earlier study using other UC MSC donors cultured in a different media formulation (Wiese et al., 2019). These data imply that Col IV synthesized by BM MSCs may only form Col IV^α121^ scaffolds, while UC MSCs synthesize the starting material for all 3 Col IV supramolecular structures and warrants further study.

These unique properties may contribute to the apparent differences in volumetric Col IV scaffolding, where the Col IV and fibronectin networks constructed by BM MSC spheroids (Fig. 3A, second row) appeared less dense than the perinatal MSC-derived matrices (Fig. 3A, bottom rows). Indeed, the highly stringent glomerular basement membrane in the kidney, which filters urine to eliminate waste while retaining proteins and other useful molecules contains two Col IV networks – a Col IV^α121^ network and a Col IV^α345^ network (Page-McCaw et al., 2025). Our gene expression data indicates that UC MSCs can synthesize both networks, as well as Col IV^α121–α556^, which would yield a more stringent Col IV network than the BM MSCs which can only produce a Col IV^α121^ network (Wiese et al., 2022).

Compaction is a natural developmental process that indicates organotypic self-organization (Baker & Chen, 2012; Domnina et al., 2020). Spontaneous compaction, as observed in this study (Fig. 1), is driven by ECM structure and stiffness (Efremov et al., 2021; Jahin et al., 2023; Kosheleva et al., 2023), actin contractility (Lee et al., 2019; Sart et al., 2014; Sharma et al., 2025; Tsai et al., 2015; Turlier & Maître, 2015), cell-matrix adhesion (Bartosh et al., 2013; Yokomine et al., 2025) and availability of matrix remodeling enzymes (van der Net et al., 2024; Yokomine et al., 2025). Here, we found that the BM MSC-derived spheroids were significantly more compact than the perinatal or adipose-derived MSC spheroids (Fig. 1), which could be the result of several key differences. The protein composition of the BM MSC matrices varied the most from the other tissue sources (Fig. 5), and they also exhibited a more open network of fibronectin and Col IV (Fig. 3A, Fig. 4A). These data suggest that the BM MSC-derived matrices may have different biomechanics than the matrices derived from the AD, UC and PL MSC donors, which we are currently assessing using biophysical techniques like atomic force microscopy and optical tweezers.

The rate of cell proliferation may also differ between MSC spheroids, leading to differences in cell density with consequent effects on compaction. A previous study showed that as the number of MSCs per spheroid increases, the packing density decreases, resulting in tightly packed, smaller spheroids and loosely packed, larger spheroids (Murphy et al., 2017). It is possible that variations in proliferation rates between spheroids, and thus differences in cell number over time, correlates with their compaction. In addition, analysis using spheroids derived from numerous donors of each sex would help discern whether different compaction rates are the result of tissue-imprinted variables or biological differences between donors irrespective of the tissue of origin.

We performed a temporal analysis of ECM deposition by MSCs cultured in multilayered sheets at different timepoints, studies that are reportedly under-represented in the literature (Fuentes et al., 2022). A previous study, using human AD MSCs cultured in DMEM/F12 plus FBS, found that matrix assembly was slower in sheet cultures than in spheroids, whereby laminin and Col IV were barely detectable in the ECM after 4 days although the MSCs stained brightly, suggesting that secretion of these proteins was just initiating (Glazieva et al., 2023). This agrees with our data, where Col IV was readily detected by ICC in matrices isolated from 4-day old spheroids (Fig. 3), when it was barely discernible in matrices from 4-day old MSC multilayers (Fig. 6A, B). Despite the relative delay in matrix assembly, the multilayered MSCs sheets established their ECM in a temporal sequence of developmental basement membrane assembly, starting with a fibronectin network (Fig. 6A, B). This foundational scaffold was then elaborated with laminin, Col IV, Col VI and additional fibronectin, recapitulating the hierarchical pattern of basement membrane assembly *in vivo* (Pompili et al., 2021). In-depth characterization of the MSC-derived matrices using proteomics is necessary to fully appreciate and compare the complement of adhesion, structural, proteoglycan, instructive and signaling components and their assembly. However, analysis of decellularized matrices is not straightforward, since conventional preparation procedures can lead to artefacts in the data, while matrisome analysis from cellularized samples is not sufficiently enriched to detect low abundance proteins (Byron et al., 2013; Diedrich et al., 2024; Frattini et al., 2024; Monteiro-Lobato et al., 2022; Naba, 2023; Ouni et al., 2020).

Environmental conditions like oxygen tension also influence matrix synthesis (Kumar et al., 2018; Lee et al., 2016; Satyam et al., 2016). For example, a recent study showed that human articular chondrocytes cultured at physiological oxygen tension (5% O_2_) produced thicker matrices with increased deposition of glycosaminoglycans and Col II as well as increased tensile strength (Dennis et al., 2020). MSC spheroids cultured under hypoxia (2% O_2_) produced more laminin, fibronectin and Col I compared to spheroids cultured at ambient (∼20%) O_2_ (Shearier et al., 2016). Since MSCs were cultivated under 5% O_2_ in our study, oxygen tension may be a key differential that contributes to the disparate outcomes between our data and other studies that maintain MSCs at ambient O_2_ tension.

Decellularized matrices derived from MSC spheroids and cell-sheet cultures represent biologically relevant scaffolds that preserve native extracellular matrix architecture, growth-factor–binding capacity, and integrin-specific adhesion cues – features that are essential for supporting cell survival and directing lineage specification in engineered tissues. Here we provide proof-of-concept that human MSCs cultured in a chemically defined bioprocess can produce basement membrane components that self-assemble into spheroid and sheet-type matrices. These naturally formed scaffolds can be effective recellularized with other cell types, demonstrating the functional integrity of the MSC-derived matrix proteins. While purified collagens, including Col I and Col IV, are known to self-assemble into matrices *in vitro* (Al-Shaer & Forde, 2025; Kotch & Raines, 2006; Yu et al., 2024), it remains unclear whether more complex matrices comprising multiple basement membrane proteins can similarly self-assemble outside the cellular context. Collectively, these findings advance the development of high fidelity, human-derived tissue models for disease modeling, drug discovery and regenerative medicine, while supporting sustainable and ethically responsible biomedical research.

## 4 Author Contributions

SNS: Collection and/or assembly of data, data analysis and interpretation, manuscript writing, critical review of the manuscript and final approval of manuscript. LRB: Conception/design, assembly of data, data analysis and interpretation, manuscript writing, critical review of the manuscript and final approval of manuscript.

## 6 Acknowledgements

The authors would like to acknowledge Tissue Regeneration Therapeutics Inc. (TRT), Toronto, Canada, for provision of human umbilical cord perivascular cells. The authors would also like to thank Braid lab members Dr. Simon Wang for his technical assistance with confocal imaging, Dr. Alaa Al-Shaer for editing the final version of the manuscript, Daniel Sloseris for characterizing decellularization efficiency and Ana Gibbert for her early optimization of the MSC spheroid cultures.

## References

Aisenbrey, E. A., & Murphy, W. L. (2020). Synthetic alternatives to Matrigel. Nature Reviews. Materials, 5(7), 539–551. 10.1038/s41578-020-0199-8

Al-Shaer, A., & Forde, N. R. (2025). Decoding collagen’s thermally induced unfolding and refolding pathways. Proceedings of the National Academy of Sciences, 122(20), e2420308122. 10.1073/pnas.2420308122

Amable, P. R., Teixeira, M. V. T., Carias, R. B. V., Granjeiro, J. M., & Borojevic, R. (2014). Protein synthesis and secretion in human mesenchymal cells derived from bone marrow, adipose tissue and Wharton’s jelly. Stem Cell Research & Therapy, 5(2), 53. 10.1186/scrt442

Baker, B. M., & Chen, C. S. (2012). Deconstructing the third dimension – how 3D culture microenvironments alter cellular cues. Journal of Cell Science, 125(13), 3015–3024. 10.1242/jcs.079509

Baker, M. (2016a). 1,500 scientists lift the lid on reproducibility. Nature, 533(7604), 452–454. 10.1038/533452a

Baker, M. (2016b). Reproducibility: Respect your cells! Nature, 537(7620), 433–435. 10.1038/537433a

Bartosh, T. J., Ylöstalo, J. H., Bazhanov, N., Kuhlman, J., & Prockop, D. J. (2013). Dynamic compaction of human mesenchymal stem/precursor cells (MSC) into spheres self-activates caspase-dependent IL1 signaling to enhance secretion of modulators of inflammation and immunity (PGE2, TSG6 and STC1). Stem Cells (Dayton, Ohio), 31(11), 10.1002/stem.1499. 10.1002/stem.1499

Bartosh, T. J., Ylöstalo, J. H., Mohammadipoor, A., Bazhanov, N., Coble, K., Claypool, K., Lee, R. H., Choi, H., & Prockop, D. J. (2010). Aggregation of human mesenchymal stromal cells (MSCs) into 3D spheroids enhances their antiinflammatory properties. Proceedings of the National Academy of Sciences, 107(31), 13724–13729. 10.1073/pnas.1008117107

Bauman, E., Feijão, T., Carvalho, D. T. O., Granja, P. L., & Barrias, C. C. (2018). Xeno-free pre-vascularized spheroids for therapeutic applications. Scientific Reports, 8(1), 230. 10.1038/s41598-017-18431-6

Byron, A., Humphries, J. D., & Humphries, M. J. (2013). Defining the extracellular matrix using proteomics. International Journal of Experimental Pathology, 94(2), 75–92. 10.1111/iep.12011

Caplan, A. I. (2017). New MSC: MSCs as pericytes are Sentinels and gatekeepers. Journal of Orthopaedic Research: Official Publication of the Orthopaedic Research Society, 35(6), 1151–1159. 10.1002/jor.23560

Catarino, C. M., Kaiser, K., Baltazar, T., Catarino, L. M., Brewer, J. R., & Karande, P. (2022). Evaluation of native and non-native biomaterials for engineering human skin tissue. Bioengineering & Translational Medicine, 7(3), e10297. 10.1002/btm2.10297

Cesarz, Z., & Tamama, K. (2016). Spheroid Culture of Mesenchymal Stem Cells. Stem Cells International, 2016, 9176357. 10.1155/2016/9176357

Cescon, M., Gattazzo, F., Chen, P., & Bonaldo, P. (2015). Collagen VI at a glance. Journal of Cell Science, 128(19), 3525–3531. 10.1242/jcs.169748

Chan, E. C., Kuo, S.-M., Kong, A. M., Morrison, W. A., Dusting, G. J., Mitchell, G. M., Lim, S. Y., & Liu, G.-S. (2016). Three Dimensional Collagen Scaffold Promotes Intrinsic Vascularisation for Tissue Engineering Applications. PLOS ONE, 11(2), e0149799. 10.1371/journal.pone.0149799

Chen, G. H., Sia, K.-C., Liu, S.-W., Kao, Y.-C., Yang, P.-C., Ho, C.-H., Huang, S.-C., Lee, P.-Y., Liang, M.-Z., Chen, L., & Huang, C.-C. (2025). Implantation of MSC spheroid-derived 3D decellularized ECM enriched with the MSC secretome ameliorates traumatic brain injury and promotes brain repair. Biomaterials, 315, 122941. 10.1016/j.biomaterials.2024.122941

Chiang, C.-E., Fang, Y.-Q., Ho, C.-T., Assunção, M., Lin, S.-J., Wang, Y.-C., Blocki, A., & Huang, C.-C. (2021). Bioactive Decellularized Extracellular Matrix Derived from 3D Stem Cell Spheroids under Macromolecular Crowding Serves as a Scaffold for Tissue Engineering. Advanced Healthcare Materials, 10(11), e2100024. 10.1002/adhm.202100024

Choi, J., Choi, W., Joo, Y., Chung, H., Kim, D., Oh, S. J., & Kim, S.-H. (2021). FGF2-primed 3D spheroids producing IL-8 promote therapeutic angiogenesis in murine hindlimb ischemia. NPJ Regenerative Medicine, 6, 48. 10.1038/s41536-021-00159-7

Corselli, M., Crisan, M., Murray, I. R., West, C. C., Scholes, J., Codrea, F., Khan, N., & Péault, B. (2013). Identification of perivascular mesenchymal stromal/stem cells by flow cytometry. Cytometry. Part A: The Journal of the International Society for Analytical Cytology, 83(8), 714–720. 10.1002/cyto.a.22313

Delabie, W., Vandewalle, V., Seghers, S., De Bleser, D., Vandekerckhove, P., & Feys, H. B. (2025). Comparison of culture media supplements identifies serum components in self-reported serum-free preparations. Stem Cell Research & Therapy, 16, 454. 10.1186/s13287-025-04561-6

Dennis, J. E., Whitney, G. A., Rai, J., Fernandes, R. J., & Kean, T. J. (2020). Physioxia Stimulates Extracellular Matrix Deposition and Increases Mechanical Properties of Human Chondrocyte-Derived Tissue-Engineered Cartilage. Frontiers in Bioengineering and Biotechnology, 8. 10.3389/fbioe.2020.590743

Diedrich, A.-M., Daneshgar, A., Tang, P., Klein, O., Mohr, A., Onwuegbuchulam, O. A., von Rueden, S., Menck, K., Bleckmann, A., Juratli, M. A., Becker, F., Sauer, I. M., Hillebrandt, K. H., Pascher, A., & Struecker, B. (2024). Proteomic analysis of decellularized mice liver and kidney extracellular matrices. Journal of Biological Engineering, 18(1), 17. 10.1186/s13036-024-00413-8

Domnina, A., Ivanova, J., Alekseenko, L., Kozhukharova, I., Borodkina, A., Pugovkina, N., Smirnova, I., Lyublinskaya, O., Fridlyanskaya, I., & Nikolsky, N. (2020). Three-Dimensional Compaction Switches Stress Response Programs and Enhances Therapeutic Efficacy of Endometrial Mesenchymal Stem/Stromal Cells. Frontiers in Cell and Developmental Biology, 8, 473. 10.3389/fcell.2020.00473

Duisit, J., Amiel, H., Wüthrich, T., Taddeo, A., Dedriche, A., Destoop, V., Pardoen, T., Bouzin, C., Joris, V., Magee, D., Vögelin, E., Harriman, D., Dessy, C., Orlando, G., Behets, C., Rieben, R., Gianello, P., & Lengelé, B. (2018). Perfusion-decellularization of human ear grafts enables ECM-based scaffolds for auricular vascularized composite tissue engineering. Acta Biomaterialia, 73, 339–354. 10.1016/j.actbio.2018.04.009

Efremov, Y. M., Zurina, I. M., Presniakova, V. S., Kosheleva, N. V., Butnaru, D. V., Svistunov, A. A., Rochev, Y. A., & Timashev, P. S. (2021). Mechanical properties of cell sheets and spheroids: The link between single cells and complex tissues. Biophysical Reviews, 13(4), 541–561. 10.1007/s12551-021-00821-w

Fonseca, V. C., Van, V., & Ip, B. C. (2023). Primary Human Cell-Derived Extracellular Matrix from Decellularized Fibroblast Microtissues with Tissue-Dependent Composition and Microstructure (p. 2023.08.15.553420). bioRxiv. 10.1101/2023.08.15.553420

Franco-Barraza, J., Beacham, D. A., Amatangelo, M. D., & Cukierman, E. (2016). Preparation of Extracellular Matrices Produced by Cultured and Primary Fibroblasts. Current Protocols in Cell Biology, 71, 10.9.1-10.9.34. 10.1002/cpcb.2

Frattini, T., Devos, H., Makridakis, M., Roubelakis, M. G., Latosinska, A., Mischak, H., Schanstra, J. P., Vlahou, A., & Saulnier-Blache, J.-S. (2024). Benefits and limits of decellularization on mass-spectrometry-based extracellular matrix proteome analysis of mouse kidney. PROTEOMICS, 24(17), 2400052. 10.1002/pmic.202400052

Fuentes, P., Torres, M. J., Arancibia, R., Aulestia, F., Vergara, M., Carrión, F., Osses, N., & Altamirano, C. (2022). Dynamic Culture of Mesenchymal Stromal/Stem Cell Spheroids and Secretion of Paracrine Factors. Frontiers in Bioengineering and Biotechnology, 10, 916229. 10.3389/fbioe.2022.916229

Glazieva, V. S., Alexandrushkina, N. A., Nimiritsky, P. P., Kulebyakina, M. A., Eremichev, R. Y., & Makarevich, P. I. (2023). Extracellular Matrix Deposition Defines the Duration of Cell Sheet Assembly from Human Adipose-Derived MSC. International Journal of Molecular Sciences, 24(23), 17050. 10.3390/ijms242317050

Gonzalez-Fernandez, T., Tenorio, A. J., Saiz, A. M., & Leach, J. K. (2022). Engineered Cell-Secreted Extracellular Matrix Modulates Cell Spheroid Mechanosensing and Amplifies Their Response to Inductive Cues for the Formation of Mineralized Tissues. Advanced Healthcare Materials, 11(10), e2102337. 10.1002/adhm.202102337

Gregory, C. A., Ma, J., & Lomeli, S. (2024). The coordinated activities of collagen VI and XII in maintenance of tissue structure, function and repair: Evidence for a physical interaction. Frontiers in Molecular Biosciences, 11. 10.3389/fmolb.2024.1376091

Hemeda, H., Giebel, B., & Wagner, W. (2014). Evaluation of human platelet lysate versus fetal bovine serum for culture of mesenchymal stromal cells. Cytotherapy, 16(2), 170–180. 10.1016/j.jcyt.2013.11.004

Hodges, G. M., & Melcher, A. H. (1976). Chemically-defined medium for growth and differentiation of mixed epithelial and connective tissues in organ culture. In Vitro, 12(6), 450–459. 10.1007/BF02806025

Huang, J., Zhang, L., Wan, D., Zhou, L., Zheng, S., Lin, S., & Qiao, Y. (2021). Extracellular matrix and its therapeutic potential for cancer treatment. Signal Transduction and Targeted Therapy, 6, 153. 10.1038/s41392-021-00544-0

Hughes, C. S., Postovit, L. M., & Lajoie, G. A. (2010). Matrigel: A complex protein mixture required for optimal growth of cell culture. Proteomics, 10(9), 1886–1890. 10.1002/pmic.200900758

Im, G.-B., Kim, S.-W., & Bhang, S. H. (2021). Fortifying the angiogenic efficacy of adipose derived stem cell spheroids using spheroid compaction. Journal of Industrial and Engineering Chemistry, 93, 228–236. 10.1016/j.jiec.2020.09.027

Jahin, I., Phillips, T., Marcotti, S., Gorey, M.-A., Cox, S., & Parsons, M. (2023). Extracellular matrix stiffness activates mechanosensitive signals but limits breast cancer cell spheroid proliferation and invasion. Frontiers in Cell and Developmental Biology, 11. 10.3389/fcell.2023.1292775

Jain, P., Rauer, S. B., Möller, M., & Singh, S. (2022). Mimicking the Natural Basement Membrane for Advanced Tissue Engineering. Biomacromolecules, 23(8), 3081–3103. 10.1021/acs.biomac.2c00402

Jayadev, R., & Sherwood, D. R. (2017). Basement membranes. Current Biology, 27(6), R207–R211. 10.1016/j.cub.2017.02.006

Jin, Y., Kim, H., Min, S., Choi, Y. S., Seo, S. J., Jeong, E., Kim, S. K., Lee, H.-A., Jo, S.-H., Park, J.-H., Park, B.-W., Sim, W.-S., Kim, J.-J., Ban, K., Kim, Y.-G., Park, H.-J., & Cho, S.-W. (2022). Three-dimensional heart extracellular matrix enhances chemically induced direct cardiac reprogramming. Science Advances, 8(50), eabn5768. 10.1126/sciadv.abn5768

Johnson, T. D., Hill, R. C., Dzieciatkowska, M., Nigam, V., Behfar, A., Christman, K. L., & Hansen, K. C. (2016). Quantification of Decellularized Human Myocardial Matrix: A Comparison of Six Patients. Proteomics. Clinical Applications, 10(1), 75–83. 10.1002/prca.201500048

Jung, S., Sen, A., Rosenberg, L., & Behie, L. A. (2010). Identification of growth and attachment factors for the serum-free isolation and expansion of human mesenchymal stromal cells. Cytotherapy, 12(5), 637–657. 10.3109/14653249.2010.495113

Kane, K. I. W., Lucumi Moreno, E., Lehr, C. M., Hachi, S., Dannert, R., Sanctuary, R., Wagner, C., Fleming, R. M. T., & Baller, J. (2018). Determination of the rheological properties of Matrigel for optimum seeding conditions in microfluidic cell cultures. AIP Advances, 8(12), 125332. 10.1063/1.5067382

Kang, S.-G., Shinojima, N., Hossain, A., Gumin, J., Yong, R. L., Colman, H., Marini, F., Andreeff, M., & Lang, F. F. (2010). Isolation and Perivascular Localization of Mesenchymal Stem Cells From Mouse Brain. Neurosurgery, 67(3), 711–720. 10.1227/01.NEU.0000377859.06219.78

Karnieli, O., Friedner, O. M., Allickson, J. G., Zhang, N., Jung, S., Fiorentini, D., Abraham, E., Eaker, S. S., Yong, tan K., Chan, A., Griffiths, S., Wehn, A. K., Oh, S., & Karnieli, O. (2017). A consensus introduction to serum replacements and serum-free media for cellular therapies. Cytotherapy, 19(2), 155–169. 10.1016/j.jcyt.2016.11.011

Keppie, S. J., Mansfield, J. C., Tang, X., Philp, C. J., Graham, H. K., Önnerfjord, P., Wall, A., McLean, C., Winlove, C. P., Sherratt, M. J., Pavlovskaya, G. E., & Vincent, T. L. (2021). Matrix-Bound Growth Factors are Released upon Cartilage Compression by an Aggrecan-Dependent Sodium Flux that is Lost in Osteoarthritis. Function, 2(5), zqab037. 10.1093/function/zqab037

Khalilgharibi, N., & Mao, Y. (2021). To form and function: On the role of basement membrane mechanics in tissue development, homeostasis and disease. Open Biology, 11(2), 200360. 10.1098/rsob.200360

Kleinman, H. K., & Martin, G. R. (2005). Matrigel: Basement membrane matrix with biological activity. Seminars in Cancer Biology, 15(5), 378–386. 10.1016/j.semcancer.2005.05.004

Klinker, M. W., Marklein, R. A., Lo Surdo, J. L., Wei, C.-H., & Bauer, S. R. (2017). Morphological features of IFN-γ-stimulated mesenchymal stromal cells predict overall immunosuppressive capacity. Proceedings of the National Academy of Sciences of the United States of America, 114(13), E2598–E2607. 10.1073/pnas.1617933114

Kosheleva, N. V., Efremov, Y. M., Koteneva, P. I., Ilina, I. V., Zurina, I. M., Bikmulina, P. Y., Shpichka, A. I., & Timashev, P. S. (2023). Building a tissue: Mesenchymal and epithelial cell spheroids mechanical properties at micro- and nanoscale. Acta Biomaterialia, Biofabrication with Spheroid and Organoid Materials, 165, 140–152. 10.1016/j.actbio.2022.09.051

Kotch, F. W., & Raines, R. T. (2006). Self-assembly of synthetic collagen triple helices. Proceedings of the National Academy of Sciences, 103(9), 3028–3033. 10.1073/pnas.0508783103

Kozlowski, M. T., Crook, C. J., & Ku, H. T. (2021). Towards organoid culture without Matrigel. Communications Biology, 4(1), 1387. 10.1038/s42003-021-02910-8

Krymchenko, R., Avila-Martinez, N., Gansevoort, M., Bakker, G.-J., Gomes, M. L. N. P., Vlig, M., Versteeg, E. M. M., Boekema, B. K. H. L., van Kuppevelt, T. H., & Daamen, W. F. (2025). Collagen-elastin dermal scaffolds enhance tissue regeneration and reduce scarring in preclinical models. Materials Today Bio, 34, 102239. 10.1016/j.mtbio.2025.102239

Kumar, P., Satyam, A., Cigognini, D., Pandit, A., & Zeugolis, D. I. (2018). Low oxygen tension and macromolecular crowding accelerate extracellular matrix deposition in human corneal fibroblast culture. Journal of Tissue Engineering and Regenerative Medicine, 12(1), 6–18. 10.1002/term.2283

Lee, J. H., Han, Y.-S., & Lee, S. H. (2016). Long-Duration Three-Dimensional Spheroid Culture Promotes Angiogenic Activities of Adipose-Derived Mesenchymal Stem Cells. Biomolecules & Therapeutics, 24(3), 260. 10.4062/biomolther.2015.146

Lee, W., Kalashnikov, N., Mok, S., Halaoui, R., Kuzmin, E., Putnam, A. J., Takayama, S., Park, M., McCaffrey, L., Zhao, R., Leask, R. L., & Moraes, C. (2019). Dispersible hydrogel force sensors reveal patterns of solid mechanical stress in multicellular spheroid cultures. Nature Communications, 10(1), 144. 10.1038/s41467-018-07967-4

Lennon, R., & Sherwood, D. R. (2025). Basement membranes at a glance. Journal of Cell Science, 138(17), jcs263947. 10.1242/jcs.263947

Li, M., Fu, T., Yang, S., Pan, L., Tang, J., Chen, M., Liang, P., Gao, Z., & Guo, L. (2021). Agarose-based spheroid culture enhanced stemness and promoted odontogenic differentiation potential of human dental follicle cells in vitro. In Vitro Cellular & Developmental Biology. Animal, 57(6), 620–630. 10.1007/s11626-021-00591-5

Liu, S., Yang, W., Li, Y., & Sun, C. (2023). Fetal bovine serum, an important factor affecting the reproducibility of cell experiments. Scientific Reports, 13, 1942. 10.1038/s41598-023-29060-7

Loomis, T., Hu, L.-Y., Wohlgemuth, R. P., Chellakudam, R. R., Muralidharan, P. D., & Smith, L. R. (2022). Matrix stiffness and architecture drive fibro-adipogenic progenitors’ activation into myofibroblasts. Scientific Reports, 12(1), 13582. 10.1038/s41598-022-17852-2

Maia, F. R., Fonseca, K. B., Rodrigues, G., Granja, P. L., & Barrias, C. C. (2014). Matrix-driven formation of mesenchymal stem cell–extracellular matrix microtissues on soft alginate hydrogels. Acta Biomaterialia, 10(7), 3197–3208. 10.1016/j.actbio.2014.02.049

Malakpour-Permlid, A., Buzzi, I., Hegardt, C., Johansson, F., & Oredsson, S. (2021). Identification of extracellular matrix proteins secreted by human dermal fibroblasts cultured in 3D electrospun scaffolds. Scientific Reports, 11(1), 6655. 10.1038/s41598-021-85742-0

Malakpour-Permlid, A., Rodriguez, M. M., Zór, K., Boisen, A., & Oredsson, S. (2025). Advancing humanized 3D tumor modeling using an open access xeno-free medium. Frontiers in Toxicology, 7. 10.3389/ftox.2025.1529360

Marinkovic, M., Tran, O. N., Block, T. J., Rakian, R., Gonzalez, A. O., Dean, D. D., Yeh, C.-K., & Chen, X.-D. (2020). Native extracellular matrix, synthesized ex vivo by bone marrow or adipose stromal cells, faithfully directs mesenchymal stem cell differentiation. Matrix Biology Plus, 8, 100044. 10.1016/j.mbplus.2020.100044

Marrella, A., Lee, T. Y., Lee, D. H., Karuthedom, S., Syla, D., Chawla, A., Khademhosseini, A., & Jang, H. L. (2018). Engineering vascularized and innervated bone biomaterials for improved skeletal tissue regeneration. Materials Today, 21(4), 362–376. 10.1016/j.mattod.2017.10.005

Minor, A. J., & Coulombe, K. L. K. (2020). Engineering a collagen matrix for cell-instructive regenerative angiogenesis. Journal of Biomedical Materials Research Part B: Applied Biomaterials, 108(6), 2407–2416. 10.1002/jbm.b.34573

Monteiro-Lobato, G. M., Russo, P. S. T., Winck, F. V., & Catalani, L. H. (2022). Proteomic Analysis of Decellularized Extracellular Matrix: Achieving a Competent Biomaterial for Osteogenesis. BioMed Research International, 2022, 6884370. 10.1155/2022/6884370

Motta, L. C. B., Pereira, V. M., Pinto, P. A. F., Mançanares, C. A. F., Pieri, N. C. G., de Olveira, V. C., Fantinato-Neto, P., & Ambrósio, C. E. (2023). 3D culture of mesenchymal stem cells from the yolk sac to generate intestinal organoid. Theriogenology, 209. 10.1016/j.theriogenology.2023.06.003

Muiznieks, L. D., & Keeley, F. W. (2013). Molecular assembly and mechanical properties of the extracellular matrix: A fibrous protein perspective. Biochimica et Biophysica Acta (BBA) - Molecular Basis of Disease, Fibrosis: Translation of Basic Research to Human Disease, 1832(7), 866–875. 10.1016/j.bbadis.2012.11.022

Murphy, K. C., Hung, B. P., Browne-Bourne, S., Zhou, D., Yeung, J., Genetos, D. C., & Leach, J. K. (2017). Measurement of oxygen tension within mesenchymal stem cell spheroids. Journal of The Royal Society Interface, 14(127), 20160851. 10.1098/rsif.2016.0851

Naba, A. (2023). Ten Years of Extracellular Matrix Proteomics: Accomplishments, Challenges, and Future Perspectives. Molecular & Cellular Proteomics: MCP, 22(4), 100528. 10.1016/j.mcpro.2023.100528

Naderi-Meshkin, H., Cornelius, V. A., Eleftheriadou, M., Potel, K. N., Setyaningsih, W. A. W., & Margariti, A. (2023). Vascular organoids: Unveiling advantages, applications, challenges, and disease modelling strategies. Stem Cell Research & Therapy, 14(1), 292. 10.1186/s13287-023-03521-2

Novoseletskaya, E., Grigorieva, O., Nimiritsky, P., Basalova, N., Eremichev, R., Milovskaya, I., Kulebyakin, K., Kulebyakina, M., Rodionov, S., Omelyanenko, N., & Efimenko, A. (2020). Mesenchymal Stromal Cell-Produced Components of Extracellular Matrix Potentiate Multipotent Stem Cell Response to Differentiation Stimuli. Frontiers in Cell and Developmental Biology, 8, 555378. 10.3389/fcell.2020.555378

Oikonomopoulos, A., van Deen, W. K., Manansala, A.-R., Lacey, P. N., Tomakili, T. A., Ziman, A., & Hommes, D. W. (2015). Optimization of human mesenchymal stem cell manufacturing: The effects of animal/xeno-free media. Scientific Reports, 5(1), 16570. 10.1038/srep16570

Olesen, K., Rodin, S., Mak, W. C., Felldin, U., Österholm, C., Tilevik, A., & Grinnemo, K.-H. (2021). Spatiotemporal Extracellular Matrix Modeling for in Situ Cell Niche Studies. Stem Cells, 39(12), 1751–1765. 10.1002/stem.3448

Oredsson, S., Malakpour-Permlid, A., Zhu, J., & Weber, T. (2025). Protocol for the use of Oredsson universal replacement medium for cell banking and routine culturing of monolayer and suspension cultures. STAR Protocols, 6(2), 103781. 10.1016/j.xpro.2025.103781

Ouni, E., Ruys, S. P. dit, Dolmans, M.-M., Herinckx, G., Vertommen, D., & Amorim, C. A. (2020). Divide-and-Conquer Matrisome Protein (DC-MaP) Strategy: An MS-Friendly Approach to Proteomic Matrisome Characterization. International Journal of Molecular Sciences, 21(23), 9141. 10.3390/ijms21239141

Page-McCaw, A., & Ferrell, N. (2025). Basement membrane structure and function: Relating biology to mechanics. Matrix Biology, 141, 16–31. 10.1016/j.matbio.2025.08.004

Page-McCaw, P. S., Pokidysheva, E. N., Darris, C. E., Chetyrkin, S., Fidler, A. L., Gallup, J., Murawala, P., Hudson, J. K., Boudko, S. P., & Hudson, B. G. (2025). Collagen IV of basement membranes: I. Origin and diversification of COL4 genes enabling metazoan multicellularity, evolution, and adaptation. The Journal of Biological Chemistry, 301(5), 108496. 10.1016/j.jbc.2025.108496

Park, J. H., Jo, S. B., Lee, J.-H., Lee, H.-H., Knowles, J. C., & Kim, H.-W. (2023). Materials and extracellular matrix rigidity highlighted in tissue damages and diseases: Implication for biomaterials design and therapeutic targets. Bioactive Materials, 20, 381–403. 10.1016/j.bioactmat.2022.06.003

Parvin Nejad, S., Lecce, M., Mirani, B., Machado Siqueira, N., Mirzaei, Z., Santerre, J. P., Davies, J. E., & Simmons, C. A. (2023). Serum- and xeno-free culture of human umbilical cord perivascular cells for pediatric heart valve tissue engineering. Stem Cell Research & Therapy, 14(1), 96. 10.1186/s13287-023-03318-3

Perugini, V., & Santin, M. (2021). A Substrate-Mimicking Basement Membrane Drives the Organization of Human Mesenchymal Stromal Cells and Endothelial Cells Into Perivascular Niche-Like Structures. Frontiers in Cell and Developmental Biology, 9, 701842. 10.3389/fcell.2021.701842

Pompili, S., Latella, G., Gaudio, E., Sferra, R., & Vetuschi, A. (2021). The Charming World of the Extracellular Matrix: A Dynamic and Protective Network of the Intestinal Wall. Frontiers in Medicine, 8. 10.3389/fmed.2021.610189

Ragelle, H., Naba, A., Larson, B. L., Zhou, F., Prijić, M., Whittaker, C. A., Del Rosario, A., Langer, R., Hynes, R. O., & Anderson, D. G. (2017). Comprehensive proteomic characterization of stem cell-derived extracellular matrices. Biomaterials, 128, 147–159. 10.1016/j.biomaterials.2017.03.008

Rakian, R., Block, T. J., Johnson, S. M., Marinkovic, M., Wu, J., Dai, Q., Dean, D. D., & Chen, X.-D. (2015). Native extracellular matrix preserves mesenchymal stem cell “stemness” and differentiation potential under serum-free culture conditions. Stem Cell Research & Therapy, 6, 235. 10.1186/s13287-015-0235-6

Ruiz, T. F. R., Ferrato, L. J., de Souza, L. G., Brito-Filho, G. E., Leonel, E. C. R., & Taboga, S. R. (2024). The elastic system: A review of elastin-related techniques and hematoxylin-eosin/phloxine applicability for normal and pathological tissue description. Acta Histochemica, 126(8), 152209. 10.1016/j.acthis.2024.152209

Sagaradze, G. D., Basalova, N. A., Efimenko, A. Y., & Tkachuk, V. A. (2020). Mesenchymal Stromal Cells as Critical Contributors to Tissue Regeneration. Frontiers in Cell and Developmental Biology, 8. 10.3389/fcell.2020.576176

Sart, S., Tsai, A.-C., Li, Y., & Ma, T. (2014). Three-dimensional aggregates of mesenchymal stem cells: Cellular mechanisms, biological properties, and applications. Tissue Engineering. Part B, Reviews, 20(5), 365–380. 10.1089/ten.TEB.2013.0537

Satyam, A., Kumar, P., Cigognini, D., Pandit, A., & Zeugolis, D. I. (2016). Low, but not too low, oxygen tension and macromolecular crowding accelerate extracellular matrix deposition in human dermal fibroblast culture. Acta Biomaterialia, 44, 221–231. 10.1016/j.actbio.2016.08.008

Schindelin, J., Arganda-Carreras, I., Frise, E., Kaynig, V., Longair, M., Pietzsch, T., Preibisch, S., Rueden, C., Saalfeld, S., Schmid, B., Tinevez, J.-Y., White, D. J., Hartenstein, V., Eliceiri, K., Tomancak, P., & Cardona, A. (2012). Fiji: An open-source platform for biological-image analysis. Nature Methods, 9(7), 676–682. 10.1038/nmeth.2019

Schneider, K. H., Goldberg, B. J., Hasturk, O., Mu, X., Dötzlhofer, M., Eder, G., Theodossiou, S., Pichelkastner, L., Riess, P., Rohringer, S., Kiss, H., Teuschl-Woller, A. H., Fitzpatrick, V., Enayati, M., Podesser, B. K., Bergmeister, H., & Kaplan, D. L. (2023). Silk fibroin, gelatin, and human placenta extracellular matrix-based composite hydrogels for 3D bioprinting and soft tissue engineering. Biomaterials Research, 27(1), 117. 10.1186/s40824-023-00431-5

Sekiguchi, R., & Yamada, K. M. (2018). Basement membranes in development and disease. Current Topics in Developmental Biology, 130, 143–191. 10.1016/bs.ctdb.2018.02.005

Sharma, S., Agashe, A., Hill, J. C., Ganguly, K., Sharma, P., Richards, T. D., Huang, W., Kaczorowski, D. J., Sanchez, P. G., Kapania, R., Phillippi, J. A., & Nain, A. S. (2025). Mechanical cues guide the formation and patterning of 3D spheroids in fibrous environments. PNAS Nexus, 4(9), pgaf263. 10.1093/pnasnexus/pgaf263

Shearier, E., Xing, Q., Qian, Z., & Zhao, F. (2016). Physiologically Low Oxygen Enhances Biomolecule Production and Stemness of Mesenchymal Stem Cell Spheroids. Tissue Engineering. Part C, Methods, 22(4), 360. 10.1089/ten.tec.2015.0465

Smith, A., Huang, M., Watkins, T., Burguin, F., Baskin, J., & Garlick, J. A. (2020). De novo production of human extracellular matrix supports increased throughput and cellular complexity in 3D skin equivalent model. Journal of Tissue Engineering and Regenerative Medicine, 14(8), 1019–1027. 10.1002/term.3071

Solomonov, I., Kollet, O., & Sagi, I. (2025). Extracellular matrix and proteolysis: Mechanisms driving irreversible changes and shaping cell behavior. The FEBS Journal. 10.1111/febs.70292

Spoerer, T. M., Larey, A. M., Asigri, W., Daga, K. R., & Marklein, R. A. (2025). High throughput morphological screening identifies chemically defined media for mesenchymal stromal cells that enhances proliferation and supports maintenance of immunomodulatory function. Stem Cell Research & Therapy, 16, 125. 10.1186/s13287-025-04206-8

Tao, F., Sayo, K., Sugimoto, K., Aoki, S., & Kojima, N. (2020). Development of a tunable method to generate various three-dimensional microstructures by replenishing macromolecules such as extracellular matrix components and polysaccharides. Scientific Reports, 10, 6567. 10.1038/s41598-020-63621-4

Trębacz, H., & Barzycka, A. (2023). Mechanical Properties and Functions of Elastin: An Overview. Biomolecules, 13(3), 574. 10.3390/biom13030574

Tsai, A.-C., Liu, Y., Yuan, X., & Ma, T. (2015). Compaction, Fusion, and Functional Activation of Three-Dimensional Human Mesenchymal Stem Cell Aggregate. Tissue Engineering. Part A, 21(9–10), 1705–1719. 10.1089/ten.tea.2014.0314

Tuftee, C., Alsberg, E., Ozbolat, I. T., & Rizwan, M. (2024). Emerging granular hydrogel bioinks to improve biological function in bioprinted constructs. Trends in Biotechnology, 42(3), 339–352. 10.1016/j.tibtech.2023.09.007

Turlier, H., & Maître, J.-L. (2015). Mechanics of tissue compaction. Seminars in Cell & Developmental Biology, 47–48, 110–117. 10.1016/j.semcdb.2015.08.001

van der Net, A., Rahman, Z., Bordoloi, A. D., Muntz, I., ten Dijke, P., Boukany, P. E., & Koenderink, G. H. (2024). EMT-related cell-matrix interactions are linked to states of cell unjamming in cancer spheroid invasion. iScience, 27(12), 111424. 10.1016/j.isci.2024.111424

Walker, C. J., Crocini, C., Ramirez, D., Killaars, A. R., Grim, J. C., Aguado, B. A., Clark, K., Allen, M. A., Dowell, R. D., Leinwand, L. A., & Anseth, K. S. (2021). Nuclear mechanosensing drives chromatin remodelling in persistently activated fibroblasts. Nature Biomedical Engineering, 5(12), 1485–1499. 10.1038/s41551-021-00709-w

Weber, T., Bajramovic, J., & Oredsson, S. (2024). Preparation of a universally usable, animal product free, defined medium for 2D and 3D culturing of normal and cancer cells. MethodsX, 12, 102592. 10.1016/j.mex.2024.102592

Wiese, D. M., & Braid, L. R. (2020). Transcriptome profiles acquired during cell expansion and licensing validate mesenchymal stromal cell lineage genes. Stem Cell Research and Therapy, 11(1), 1–7. 10.1186/S13287-020-01873-7/FIGURES/3

Wiese, D. M., Ruttan, C. C., Wood, C. A., Ford, B. N., & Braid, L. R. (2019). Accumulating Transcriptome Drift Precedes Cell Aging in Human Umbilical Cord-Derived Mesenchymal Stromal Cells Serially Cultured to Replicative Senescence. STEM CELLS Translational Medicine, 8(9), 945–958. 10.1002/SCTM.18-0246

Wiese, D. M., Wood, C. A., Ford, B. N., & Braid, L. R. (2022). Cytokine Activation Reveals Tissue-Imprinted Gene Profiles of Mesenchymal Stromal Cells. Frontiers in Immunology, 13. https://www.frontiersin.org/articles/10.3389/fimmu.2022.917790

Xu, J., Lian, W., Chen, J., Li, W., Li, L., & Huang, Z. (2020). Chemical-defined medium supporting the expansion of human mesenchymal stem cells. Stem Cell Research & Therapy, 11, 125. 10.1186/s13287-020-01641-7

Yin, S., Wu, H., Huang, Y., Lu, C., Cui, J., Li, Y., Xue, B., Wu, J., Jiang, C., Gu, X., Wang, W., & Cao, Y. (2024). Structurally and mechanically tuned macroporous hydrogels for scalable mesenchymal stem cell–extracellular matrix spheroid production. Proceedings of the National Academy of Sciences, 121(28), e2404210121. 10.1073/pnas.2404210121

Yokomine, S., Makino, T., Nagao, E., & Nakazawa, K. (2025). Floating or adherent hepatocyte spheroid cultures using microwell chips with polyethylene glycol or polyimide surfaces. Biomedical Materials, 20(3), 035009. 10.1088/1748-605X/adc17d

Yu, L. T., Kreutzberger, M. A. B., Bui, T. H., Hancu, M. C., Farsheed, A. C., Egelman, E. H., & Hartgerink, J. D. (2024). Exploration of the hierarchical assembly space of collagen-like peptides beyond the triple helix. Nature Communications, 15(1), 10385. 10.1038/s41467-024-54560-z

Yurchenco, P. D. (2011). Basement Membranes: Cell Scaffoldings and Signaling Platforms. Cold Spring Harbor Perspectives in Biology, 3(2), a004911. 10.1101/cshperspect.a004911

Zhang, X., Battiston, K. G., Labow, R. S., Simmons, C. A., & Santerre, J. P. (2017). Generating favorable growth factor and protease release profiles to enable extracellular matrix accumulation within an *in vitro* tissue engineering environment. Acta Biomaterialia, 54, 81–94. 10.1016/j.actbio.2017.02.041

Zhang, Y., Feng, Y., Shao, Q., Jiang, Z., & Yang, G. (2023). Rapid formation of 3D: Decellularized extracellular matrix spheroids for enhancing bone formation. Journal of Biomedical Materials Research. Part A, 111(3), 378–388. 10.1002/jbm.a.37471

Zhao, W., Zhu, J., Hang, J., & Zeng, W. (2022). Biomaterials to promote vascularization in tissue engineering organs and ischemic fibrotic diseases. MedComm – Biomaterials and Applications, 1(1), e16. 10.1002/mba2.16

Zhou, Y., Chen, H., Li, H., & Wu, Y. (2017). 3D culture increases pluripotent gene expression in mesenchymal stem cells through relaxation of cytoskeleton tension. Journal of Cellular and Molecular Medicine, 21(6), 1073–1084. 10.1111/jcmm.12946

